# An intranasal nanoparticle STING agonist has broad protective immunity against respiratory viruses and variants

**DOI:** 10.1101/2022.04.18.488695

**Authors:** Ankita Leekha, Arash Saeedi, Monish Kumar, Samiur Rahman Sefat, Melisa Martinez-Paniagua, Mohsen Fathi, Rohan Kulkarni, Sujit Biswas, Daphne Tsitoura, Xinli Liu, Laurence J.N. Cooper, Manu Sebastian, Brett L. Hurst, Navin Varadarajan

**Author notes:** To whom correspondence should be addressed: Navin Varadarajan, Department of Chemical and Biomolecular Engineering, University of Houston, Houston, Texas TX 77204, USA.

## Abstract

Respiratory viral infections, especially Influenza (endemic) or SARS-CoV-2 (pandemic since 2020), cause morbidity and mortality worldwide. Despite remarkable progress in the development and deployment of vaccines, they are clearly impacted by the rapid emergence of viral variants. The development of an off-the-shelf, effective, safe, and low-cost drug for prophylaxis against respiratory viral infections is a major unmet medical need. Here, we developed NanoSTING, a liposomally encapsulated formulation of the endogenous STING agonist, 2’-3’ cGAMP, to function as an immunoantiviral. NanoSTING rapidly activates the body’s innate immune system to facilitate a broad-spectrum antiviral response against SARS-CoV-2 and influenza variants in hamsters and mice. We demonstrate that a single intranasal dose of NanoSTING can: (1) treat infections throughout the respiratory system and minimize clinical symptoms, (2) protect against highly pathogenic strains of SARS-CoV-2 (alpha and delta), (3) provide durable protection against reinfection from the same strains without the need for retreatment, (4) prevent transmission of the highly infectious SARS-CoV-2 Omicron strain, and (5) provide protection against both oseltamivir-sensitive and resistant strains of influenza. Mechanistically, administration of NanoSTING rapidly upregulated interferon-stimulated and antiviral pathways in both the nasal turbinates and lung. Our results support using NanoSTING as a thermostable, immunoantiviral with broad-spectrum antiviral properties making it appealing as a therapeutic for prophylactic or early post-exposure treatment.

## Introduction

Within the last 20 years, we experienced four global respiratory epidemics/pandemics: severe acute respiratory syndrome (SARS) in 2003, influenza H1N1 in 2009, Middle East respiratory syndrome coronavirus in 2012, and severe acute respiratory syndrome coronavirus 2 (SARS-CoV-2) in 2019. These pandemics added to the global burden of existing threats like seasonal Influenza and respiratory syncytial virus (RSV)^1, 2^. The recent COVID-19 pandemic caused by SARS-CoV-2 has led to 6.17 million deaths (April 2022), while current outbreaks driven by new variants of concern (VOC’s) continue to be reported worldwide. Three classes of interventions comprise the modern arsenal of responses against respiratory viruses; vaccines, antibodies, and antivirals^3–5^. All these three interventions require significant time for identification and characterization of the virus, development, and rapid testing to identify the emerging pathogen, followed by manufacturing and global distribution of therapeutics or vaccines. With regards to rapidly mutating viruses such as the RNA viruses, all these three modalities are prone to failure due to the high mutation rate of the virus coupled with insufficient and ineffective protection, which facilitates the evolution of resistant variants^6–8^.

Respiratory viruses enter the body and initiate replication in the respiratory tract. In response to the initial infection, the host elicits a multi-faceted innate immune response, typically characterized by the antiviral interferon (IFN) response, and the ensuing battle between the host immune system and the virus dictates the progression and outcome of infection^9, 10^. Despite the IFN antagonistic mechanisms evolved by pathogens, these innate immunity responses dominate in most individuals and result in primarily asymptomatic infection or localized in the airways illness, still permitting an onward transmission of virus^11, 12^. If, however, the host’s innate immune response is suboptimal for any reason, including genetic defects or autoantibodies against IFNs, the viral infection progresses, leading to disseminated disease and even mortality^13, 14^. Ensuring robust antiviral innate immune responses in the airways is central to controlling viral infection, replication, transmission, and disease outcomes. Although conceptually straightforward, harnessing this host antiviral response is challenging. Direct administration of IFN proteins in clinical trials for COVID-19 has yielded mixed results with undesirable side effects^15, 16^. It is thus clear that the location, duration, and timing of host-directed immunotherapies are necessary to ensure the activation of the appropriate antiviral pathways that balance efficacy without causing tissue damage and toxicity.

The stimulator of the interferon genes (STING) pathway is an evolutionary-conserved cellular sensor of cytosolic double-stranded DNA (dsDNA), enabling a broad innate immune response against viruses^17, 18^. Mechanistically, activation of STING fosters an antiviral response that involves not just the type I and III interferons (IFN-I and IFN-III) but also additional pathways independent of interferon signaling^19, 20^. In humans, pre-activated STING mediated immunity in the upper airways controls early SARS-CoV-2 infection in children and can explain why children are much less susceptible to advanced disease^21, 22^. Multiple reports have demonstrated that supra-physiologic activation of STING inhibits replication of viruses, including coronaviruses and that viruses have evolved mechanisms to prevent the optimal activation of STING within the host^22, 23^.

Here, we demonstrate that intranasal delivery of a nanoparticle formulation of cyclic guanosine monophosphate–adenosine monophosphate (cGAMP), termed NanoSTING, enables the sustained release of cGAMP to both the nasal compartment and the lung for up to 48 h. The delivered cGAMP activates multiple antiviral pathways and facilitates IFN-I and IFN-III mediated responses. We deployed quantitative modeling based upon SARS-CoV-2 infections in humans to confirm that pre-exposure or early post-exposure prophylaxis with even small amounts of NanoSTING has high translational potential. In addition, we tested the ability of NanoSTING to protect against multiple VOCs of SARS-CoV-2 in hamsters and multiple variants of influenza A in mice. In these animal models, NanoSTING treatment prevented clinical disease, reduced viral titers by several orders of magnitude, reduced transmission, and enabled durable protection from reinfection. The stability, ease of administration, and the comprehensive nature of the immune response elicited make NanoSTING an ideal intranasal broad-spectrum antiviral independent of the type of respiratory virus and variants.

## Results

### Preparation, Characterization, and Stability of NanoSTING

NanoSTING is a liposomal formulation optimized for the delivery of cGAMP to the respiratory tract (**Figure 1A**). The nanoparticles promote stability and have been shown to promote delivery to alveolar macrophages, facilitating the initiation of innate immune responses in the upper airways and the lung^24, 25^. Dynamic light scattering (DLS) analysis revealed that the mean particle diameter of NanoSTING was 98 nm, with a polydispersity index of 25.1 % (**Figure S1A**). The zeta potential of NanoSTING was -40 mV (**Figure S1B**). We confirmed the ability of NanoSTING to induce interferon responses by using THP-1 monocytic cells modified to conditionally secrete luciferase downstream of an interferon regulatory factor (IRF) responsive promoter (**Figure 1B**). We stimulated THP-1 dual cells with NanoSTING at a dose from 2.5-10.0 µg and performed kinetic measurements for 24 h by measuring the luciferase activity in the supernatant. We observed a low level of luciferase activity at 6 h, and secretion was maximal at 24 h with 5 µg and 10 µg NanoSTING (**Figure 1C**). We next systematically measured the stability of the nanoparticles by assessing particle sizes and zeta potential of NanoSTING at two different temperatures, 25°C and 37°C. While the hydrodynamic diameter of NanoSTING was essentially unchanged at 25 °C over a period of 30 days (**Figure 1D**), there was a slight increase in hydrodynamic diameter at 37°C after 2 weeks (mean: 114 nm at 25°C and 154 nm at 37 °C) [**Figure 1E**]. We did not observe a change in zeta potential at both temperatures (−45mV at 25 °C and 37°C) [**Table S1**]. These results demonstrate that NanoSTING is immunologically active and that the nanoparticle remains stable even without refrigeration.

**Figure 1:**
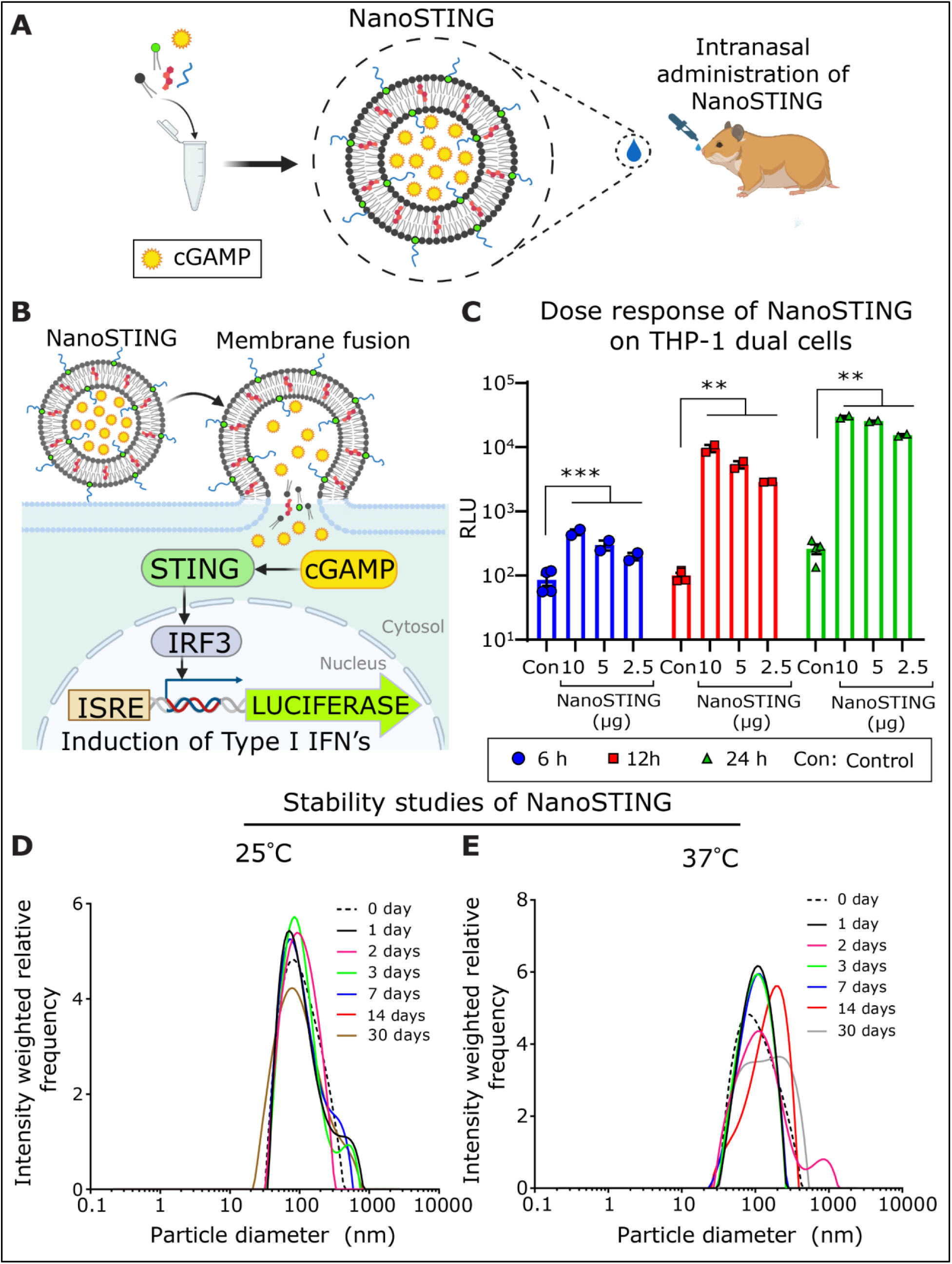
Synthesis, characterization, and stability of NanoSTING. (A) Overall schematic of the formulation and intranasal delivery of NanoSTING to animals. (B) Mechanisms of NanoSTING delivery and signaling pathways. THP1-dual cells stably express the secreted form of luciferase under the control of a synthetic interferon responsive promoter. Activation of the IRF pathway leads to the secretion of luciferase in the cell culture supernatant. (C) Kinetics of the induction of luciferase in THP1-dual cells by varying concentrations of the NanoSTING. RLU: relative light units. (D, E) Distribution of NanoSTING liposomal particle sizes at 25°C and 37°C measured by dynamic light scattering (DLS). See also Figure S1 and Table S1 *Analysis was performed using a Mann-Whitney test. Vertical bars show mean values with error bar representing SEM. Mann-Whitney test: ****p < 0.0001; ***p < 0.001; **p < 0.01; *p < 0.05; ns: not significant*.

### NanoSTING delivers cGAMP across mucosa, leading to sustained Interferon-beta (IFN-β) secretion in the nasal compartment

Although cGAMP is a potent natural activator of STING and therefore acts as an immunotransmitter, its clinical utility is hampered by lack of cellular penetration and rapid degradation by plasma ectonucleotide pyrophosphatase phosphodiesterase 1 (ENPP1), leading to an *in vivo* half-life of only ∼35 min^26^. We first characterized the ability of NanoSTING to mediate the delivery of cGAMP in the nasal compartment of mice. We delivered varying amounts of NanoSTING (10-40 μg) intranasally to groups of BALB/c mice, harvested the nasal turbinates and lungs, and assayed cGAMP using quantitative ELISA (**Figure 2A**). We observed a dose-dependent increase in the concentration of cGAMP in the nasal turbinates; at the low dose (10 μg), we quantified cGAMP up to 12 h with a return to baseline at 24 h, whereas at the higher doses (20-40 μg), we detected cGAMP for 24 h with a return to baseline at 48 h (**Figure 2B**). In the lungs, cGAMP was only detectable at the higher concentrations (20 and 40 μg) [**Figure 2C]**. We also profiled the sera of these same animals and observed that cGAMP was not detected at any timepoints in circulation, even at the highest dose (40 μg) [**Figure S2**]. These data confirmed that NanoSTING can transport cGAMP to the cells of the nasal passage in a concentration and time-dependent manner without systemic exposure.

**Figure 2:**
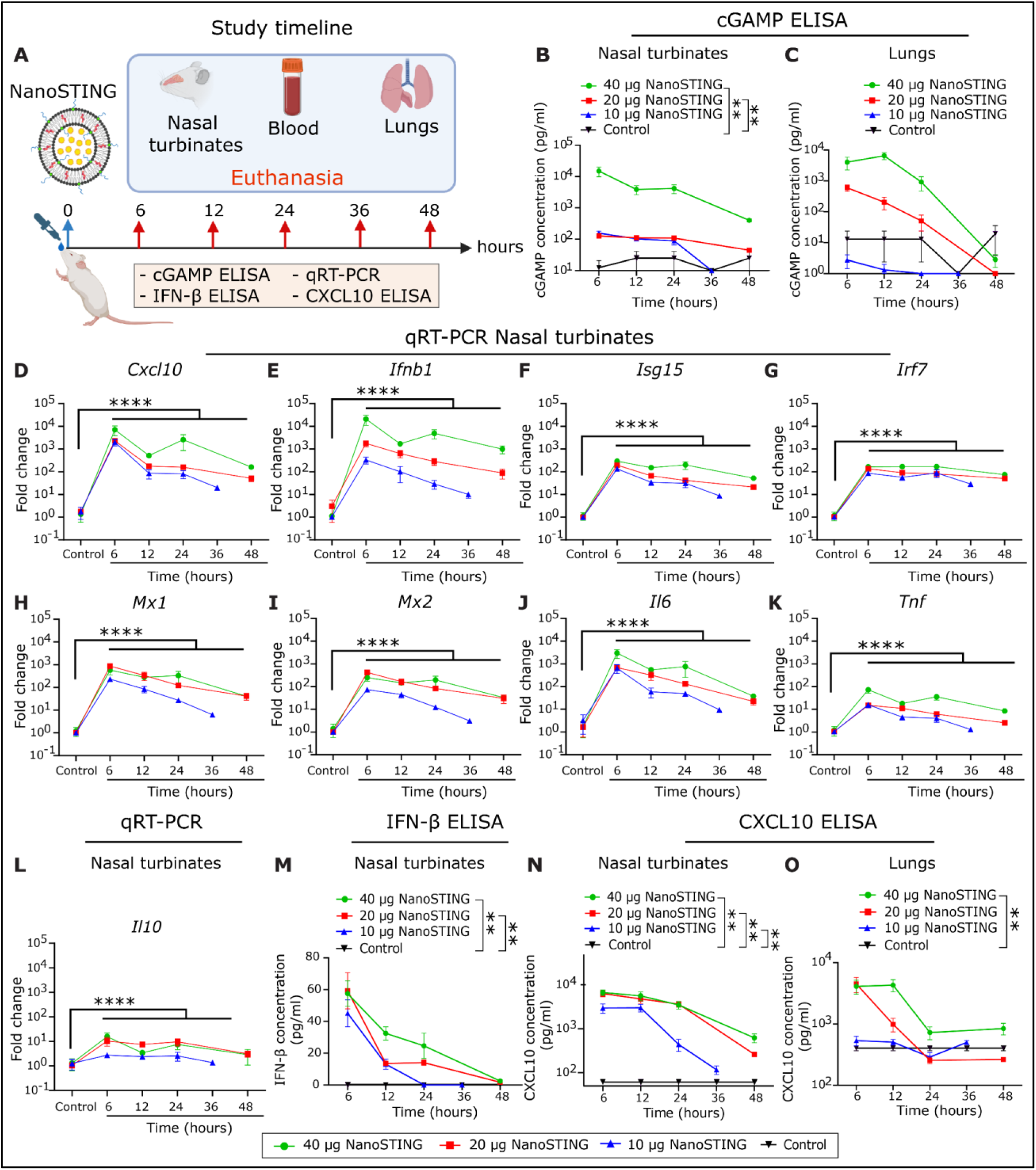
Pharmacokinetic and pharmacodynamics profiling of NanoSTING reveals prolonged delivery of cGAMP and induction of ISGs in the nasal compartment of mice. (A) Groups of 3-4 BALB/c mice were treated with single doses of NanoSTING (10 µg, 20 µg, or 40 µg), and euthanized subsets at 6 h, 12 h, 24 h, 36 h, and 48 h followed by collection of blood, nasal turbinates, and lungs. cGAMP ELISA, IFN-β ELISA, CXCL10 ELISA, and qRT-PCR (nasal turbinates) were used as the primary endpoints. (B, C) ELISA quantification of cGAMP in the nasal turbinates and lungs of mice after treatment with NanoSTING. (D-L) Fold change in gene expression for NanoSTING treated (40 µg in green, 20 µg in red, and 10 µg in blue) mice and control mice was quantified in RNA extracted from nasal turbinates by qRT-PCR (Primer sequences are provided in Table S2). (M) Quantification of IFN-β concentration in mouse nasal tissue using quantitative ELISA. (N, O) Quantification of CXCL10 levels in mouse nasal tissue and lungs using quantitative ELISA. See also Figures S2 and S3 *For ELISA and fold changes in gene expression, analysis was performed using a Mann-Whitney test. Mann-Whitney test: ****p < 0.0001; ***p < 0.001; **p < 0.01; *p < 0.05; ns: not significant*.

The biological implications of NanoSTING’s ability to deliver cGAMP and thus activate the STING pathway were evaluated using a panel of 10 genes to comprehensively measure the immune response. The panel comprised of the effector cytokines, C–X–C motif chemokine ligand 10 (*Cxcl10*) and interferon beta (*Ifnb*); Interferon stimulated genes (ISG) including *Isg15*, Interferon regulatory factor 7 (*Irf7*), myxovirus resistance proteins 1 & 2 (*Mx1* and *Mx2*), and Interferon-induced protein with tetratricopeptide repeats 1 (*Ifit1*); and non-specific pro-inflammatory cytokines (*Il6*, *Il10*, and *Tnf*). BALB/c mice received varying doses of intranasal NanoSTING, and quantitative qRT-PCR was performed on the nasal turbinates (6-48 h) [**Figure 2A**]. The effector cytokines *Cxcl10* and *Ifnb* showed maximal induction (7,000 to 20,000-fold induction) that remained elevated at 48 h (**Figures 2D-E**). The five ISGs demonstrated strong induction from 6 h (300 to 1,000-fold) to 24 h, followed by decline from 24-48 h (**Figures 2F-I and S3**). NanoSTING’s inflammatory response was linked to the IFN pathway as the pro-inflammatory cytokine *Il6* showed brief induction at 6 h (5,000-fold), declined significantly by 24 h, and returned to baseline levels at 48 h (**Figure 2J**). Furthermore, *Tnf* and *Il10* showed only weak induction (15 to 60-fold) [**Figures 2K-L**]. These results demonstrate that NanoSTING elicits a rapid and sustained inflammatory response triggering both effector cytokines and ISGs, but only minimal activation of non-specific pro-inflammatory cytokines.

Since the qRT-PCR data suggested strong induction of the effector cytokines, *Ifnb* and *Cxcl10*, we quantified the concentration of IFN-β and CXCL10 proteins in the nasal turbinates. Consistent with the transcriptional data, quantitative ELISA confirmed that both IFN-β and CXCL10 could be detected in the nasal turbinates and lungs for up to 24 h (**Figures 2M-O**). We also tested the serum of these same animals, but we did not observe neither IFN-β nor CXCL10 (**Figure S2**), confirming that stimulation of innate immunity by intranasal NanoSTING was localized in the airways, without associated induction of systemic pro-inflammatory activity.

### RNA-sequencing confirms a robust IFN-I signature in the lungs of hamsters following intranasal NanoSTING administration

We next wanted to investigate the impact of intranasal NanoSTING administration on the lungs of Syrian Golden hamsters (*Mesocricetus auratus*). The hamster is a well-characterized model for the SARS-CoV-2 challenge and mimics severe disease in humans; animals demonstrate easily quantifiable clinical disease characterized by rapid weight loss, very high viral titers in the lungs, and extensive lung pathology^27^. Additionally, unlike the K18-hACE2 transgenic model, hamsters recover from the disease (like most humans) and hence offer the opportunity to study the impact of treatments in the disease process, as well as in virus transmission^27, 28^.

We studied biodistribution by altering the transport volume of intranasally delivered NanoSTING. It has been previously demonstrated that lower volumes lead to more efficient delivery to the nasal passage while larger volumes facilitate delivery to the lung^29^. Intranasal administration of Evan’s blue dye in low and high volumes (40 μL and 120 μL) resulted in staining of the nasal turbinates, lungs, and stomach in hamsters (**Figures S4C-E)**. However, at both volumes, there was a significant amount of the dye delivered to the nasal turbinates and lung (intended target organs) [**Figure S4C**], and the normalized ratio of distribution to these tissues was independent of the volume of administration (**Figures S4D-E**). These results suggested that biodistribution to the lung/nasal compartments after intranasal delivery of liquid formulations is not impacted by the volume of inoculum in the hamster model.

To assess the impact of intranasal NanoSTING on the lung, we administered one group (n=4/group) of hamsters with daily doses of NanoSTING (60 µg) for four consecutive days. We used naive hamsters as controls (n=4/group). Both groups of hamsters showed no differences in clinical signs, such as temperature or bodyweight (**Figures S4A-B**). On day 5, we isolated the lungs from hamsters for unbiased whole-transcriptome profiling using RNA-sequencing (RNA-seq). At a false-discovery rate (FDR *q-value* < 0.25), we identified a total of 2,922 differentially expressed genes (DEGs) between the two groups (**Figure 3A**). A type I IFN response was induced in NanoSTING treated lungs, comprising canonical ISGs, including *Mx1*, *Isg15*, *Uba7*, *Ifit2*, *Ifit3*, *Ifit35*, *Irf7*, *Adar*, and *Oas2* (**Figure 3B**)^30^. The effector cytokines, *Cxcl9-11* and *Ifnb,* were also induced in treated hamsters (**Figure 3C**), as were the direct antiviral proteins such as *Ddx60* and *Gadd45g* (**Figure 3D**)^31, 32^. We performed gene-set enrichment analysis (GSEA) to compare the differentially induced pathways upon treatment with NanoSTING. We interrogated the changes in these populations against the Molecular Signatures Database (Hallmark, C2, and C7 curated gene sets). We observed a distinct cluster of pathways related to both type I and type III interferons in the lungs of NanoSTING treated mice. We confirmed the specificity of the response by qRT-PCR analyses by quantifying *Mx1-2*, *Isg15*, *Irf7*, *Cxcl11*, *Ifnb*, *Il6*, and *Il10* (**Figure S5**). Since the gene signature of interferon-independent activities of STING is known^19^, we performed GSEA and confirmed that NanoSTING activates interferon-independent pathways (**Figures 3E-F)**. In aggregate, these results demonstrate that cGAMP mediated activation of STING by NanoSTING efficiently engages both interferon-dependent and interferon-independent antiviral pathways in the lung.

**Figure 3:**
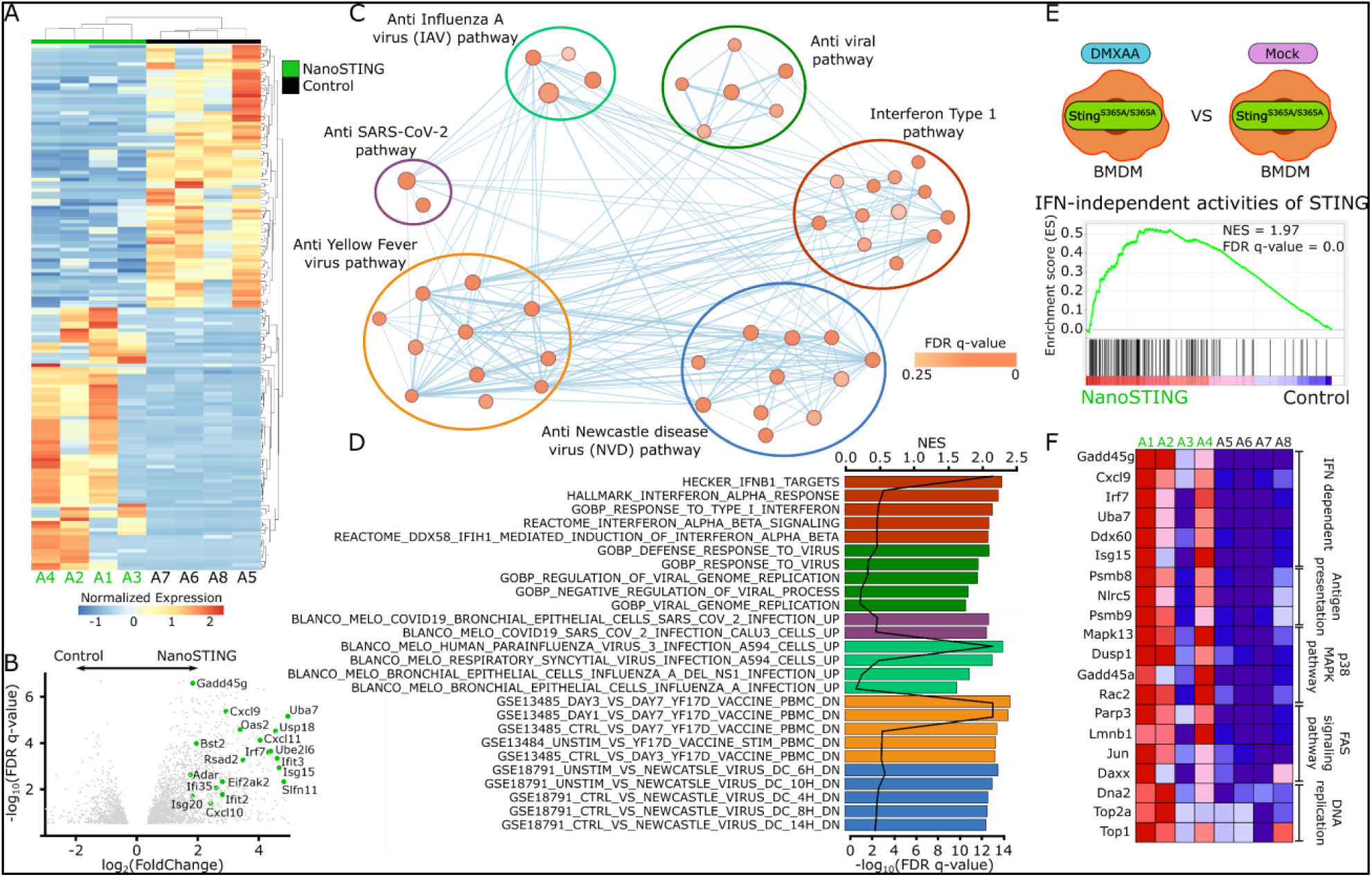
RNA-sequencing identifies the activation of IFN-dependent and IFN-independent pathways in the lungs of hamsters treated with NanoSTING. (A) Heatmap of top 50 differentially expressed genes (DEGs) between NanoSTING treated lungs (marked as green) and control lungs (marked as black). (B) The volcano plots of DEGs comparing NanoSTING treated and control animals. (C) Geneset enrichment analyses (GSEA) of C2 and C7 curated pathways visualized using Cytoscape. Nodes (red and blue circles) represent pathways, and the edges (blue lines) represent overlapping genes among pathways. The size of nodes represents the number of genes enriched within the pathway, and the thickness of edges represents the number of overlapping genes. The color of nodes was adjusted to an FDR q value ranging from 0 to 0.25. Clusters of pathways are labeled as groups with a similar theme. (D) The normalized enrichment score (NES) and false-discovery rate (FDR) *q values* of top antiviral pathways curated by GSEA analysis. (E) GSEA of IFN-independent activities of STING pathway activated in the lung of NanoSTING treated animals. The schematic represents the comparison that was made between samples collected from GSE149744 dataset to generate the pathway gene set. (F) The expression of genes in lungs associated with IFN-dependent and IFN-independent antiviral pathways between NanoSTING and control groups. See also Figures S4, S5, and Table S3

### Quantitative modeling predicts that early treatment with NanoSTING will dampen viral replication

The *in vivo* mechanistic experiments demonstrated that NanoSTING induces a broad antiviral response by engaging the innate immune system. To investigate potential efficacy, we used a mathematical model in combination with human viral titer data to identify the treatment window and quantify the relative amount of type I IFN (or related pathways) elicited by NanoSTING required for therapeutic benefit^33, 34^. To simplify the framework of the model, we assumed that *in vivo* cGAMP only works to stimulate interferon responses. With this assumption, we modeled the range of relative interferon ratios (RIR, 0-1) we need to elicit via NanoSTING in comparison to the population level peak interferon responses observed upon SARS-CoV-2 infection (**Figures 4A-B)** [See *supplementary methods*], and investigated the influence on viral elimination. Based on the model, an RIR of just 0.27 (27% of natural infection) would be sufficient to achieve a 50% reduction in viral titer (based on the area under the curve, AUC), and RIR values of at least 0.67 will achieve 100% reduction in viral titers (**Figure 4C**). We next modeled the window of initiation of treatment which revealed that intervention would be most effective when initiated within 2 days after infection (**Figure 4D**). By contrast, if the treatment is initiated after the peak of viral replication, even with an RIR of 1, improvement in outcomes cannot be readily realized (**Figures 4D and S6C**). Collectively, these results from quantitative modeling predicted that: (1) a single dose of NanoSTING is adequate to elicit only a moderate amount of IFN, and this is likely within reach since our data supports large induction of IFN-β (**Figures 2E, 2M, and 3F**) and since natural infection with viruses like SARS-CoV-2 and Influenza A is known to suppress interferon production^35–37^, and (2) the optimal treatment window was both as prophylaxis and soon after infection.

**Figure 4.**
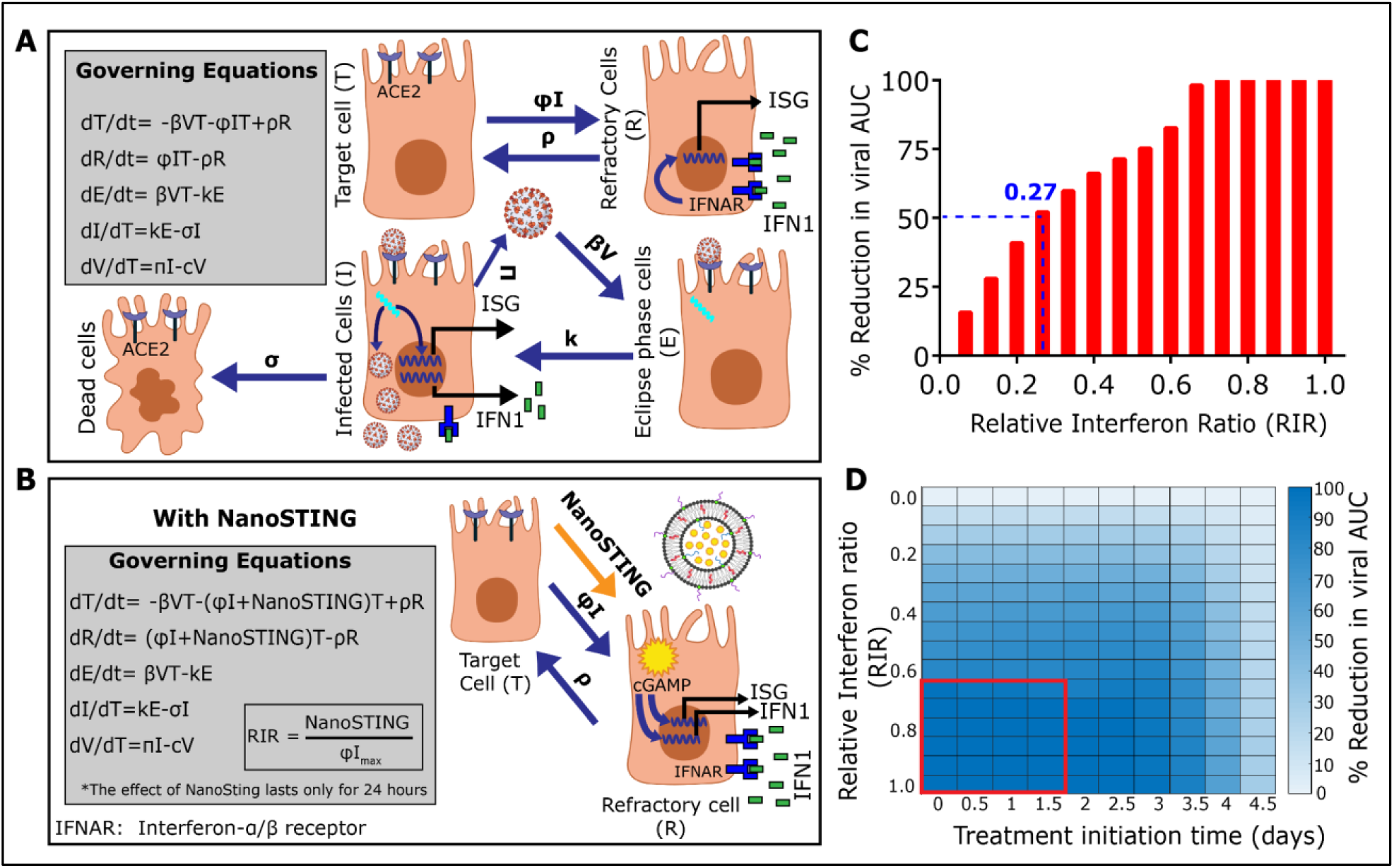
Quantitative modeling of the dynamics of replication of SARS-CoV-2. (A, B) Schematic representing rate constants and equations governing viral dynamics during (A) natural infection and (B) in the presence of NanoSTING treatment. (C) Reduction in the viral area under the curve (AUC) at different NanoSTING efficacies (RIR) compared to natural infection. The treatment is initiated on day 0, and we assume that the effects of NanoSTING treatment only last for 24 h. (D) Heatmap of viral AUC with varying NanoSTING efficacy and treatment initiation time. The red box represents the combination with close to 100% reduction in viral AUC. See also Figure S6

### NanoSTING protects against the SARS-CoV-2 Delta VOC

Based on the prediction of the modeling studies, we evaluated if a single dose of NanoSTING protects hamsters from SARS-CoV-2 infection. The Delta VOC (B.1.617.2) was chosen because it causes both upper and lower tract disease and has increased disease severity compared with prior VOCs (Wuhan and Beta strains).^38^ We treated groups of 12 Syrian golden hamsters with a single intranasal dose of 120 µg NanoSTING, and 24 h later, we infected the hamsters with ∼3 x 10^4^ 50% Cell culture infectious dose (CCID50) of the Delta VOC through the intranasal route (**Figure 5A**). In the placebo-treated (PBS) group, hamsters exhibited weight loss, with a mean peak weight loss of 8.3 %. By contrast, hamsters treated with NanoSTING were largely protected from weight loss (mean peak weight loss of 2.0 %) (**Figure 5 C**). This small amount of weight loss in hamsters is similar to the results obtained by adenoviral vectored vaccines challenged with either the Wuhan or Beta strains^39^. We quantified the infectious viral titers by sacrificing six hamsters on day 2. Even with the highly infectious Delta strain, NanoSTING reduced infectious viral titers in the lung post 2 days of infection by 300-fold compared to placebo-treated animals (**Figure 5D**). This reduction in viral titers in the lung closely correlates with the prevention of weight loss in these animals and models protection similar to clinical human disease. We also quantified the viral titers in the nasal compartment and observed that treatment with NanoSTING reduced infectious viral titers in the nasal compartment 2 days post infection by 1,000-fold compared to placebo-treated animals (**Figure 5E**). The reduction in viral replication in the nasal compartment models propensity of human transmission and confirms that treatment with NanoSTING decreases the likelihood of transmission. To map the duration of efficacy of prophylactic NanoSTING treatment, we administered a single intranasal dose of NanoSTING (120 µg) and challenged the hamsters 72 h later with ∼3 x 10^4^ CCID50 of the Delta VOC (**Figure S7A**). Even when administered at 72 h before exposure, NanoSTING showed moderate protection from weight loss and a significant reduction in infectious viral titers (**Figures S7B-D**). Since our mathematical model also predicts that NanoSTING can be used to control infection after viral exposure, we tested the efficacy of intranasal NanoSTING delivered 6 h after exposure to the Delta VOC (**Figure S8A**). We observed 340-fold and 13-fold reduction in infectious virus in the nasal passage and lung, respectively (**Figures S8B-C**). These results show that a single dose treatment with NanoSTING can effectively minimize clinical symptoms, protect the lung, and reduce infectious viruses in the nasal passage.

**Figure 5:**
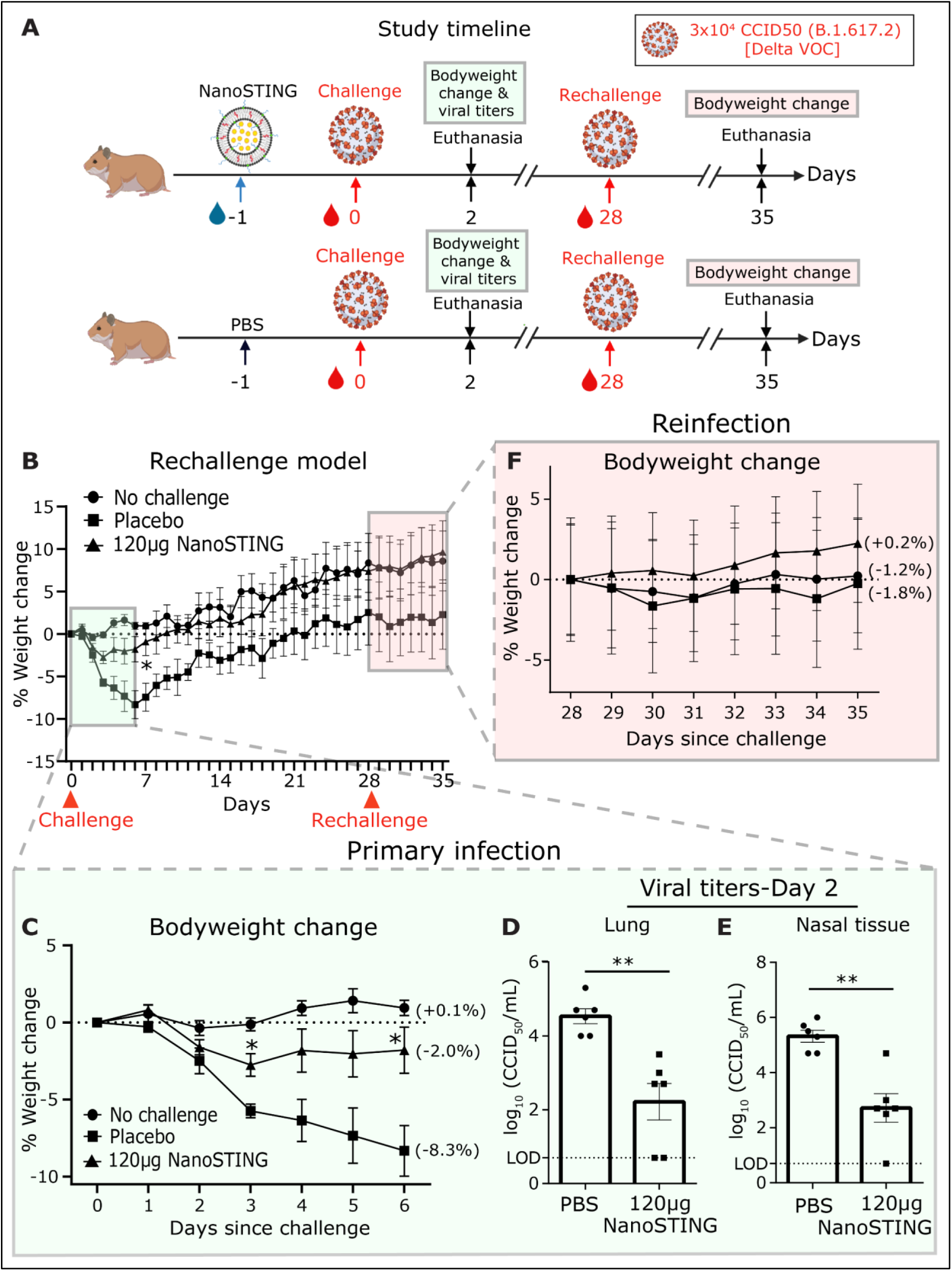
Treatment with NanoSTING protects hamsters during the primary challenge and prevents reinfection with the pathogenic SARS-CoV-2 Delta (B.1.617.2) VOC. (A) We treated groups of 12 hamsters, each with a single dose of 120 µg NanoSTING and later challenged with ∼3 x 10^4^ CCID50 of SARS-CoV-2 Delta VOC on day 0 by the intranasal route. We euthanized half of the hamsters (n=6) hamsters on day 2 and determined viral titers of lung and nasal tissues. We rechallenged the remaining 6 hamsters on day 28 and tracked the bodyweight change until day 35. (B) Percent bodyweight change compared to the baseline at the indicated time intervals. (C) Percent bodyweight change monitored during the primary infection (day 0-day 6). (D, E) Viral titers measured by plaque assay in nasal tissues and lungs post day 2 of infection. The dotted line indicates limit of detection of the assay (LOD). (F) Percent bodyweight change monitored after rechallenge (day 28-day 35). See also Figures S7 and S8 *For viral titers, analysis was performed using a Mann-Whitney test. Vertical bars show mean values with error bar representing SEM. Each dot represents an individual hamster. Weight data was compared via mixed-effects model for repeated measures analysis. Lines depict group mean body weight change from day 0; error bars represent SEM. Asterisks indicate significance compared to the placebo-treated animals at each time point*. *\*\*\*\*p < 0.0001; ***p < 0.001; **p < 0.01; *p < 0.05; ns: not significant*.

### Treatment with NanoSTING induces protection against SARS-CoV-2 reinfection

One of the advantages of enhancing the innate immunity to clear a viral infection is that this process mimics natural host defense and minimizes the danger of clinical symptoms. We hypothesized that NanoSTING treatment via activation of the innate immune system also facilitates immunological memory against reinfection, without the need for additional treatment. To test this hypothesis, we intranasally treated hamsters (n=12/group) with NanoSTING (120 µg), and 24 h later challenged with ∼3 x 10^4^ of the Delta VOC (**Figure 5A**). On day 28, we rechallenged the hamsters with the Delta VOC. The untreated animals suffered significant weight loss during the primary challenge but were largely protected during the secondary challenge (**Figure 5F**). By contrast, NanoSTING treated hamsters showed minimal weight loss during the primary challenge, and this did not compromise immunological memory. Indeed, NanoSTING treated hamsters were completely protected from weight loss during the secondary challenge, and their bodyweight was identical to animals that were not previously challenged (**Figure 5F**). These results suggest that a single intranasal treatment with NanoSTING activates the antiviral program of innate immunity, preventing clinical disease during primary infection while offering durable protection from reinfection.

### NanoSTING treatment protects against the SARS-CoV-2 Alpha VOC

We next evaluated the impact of treatment with varying doses of NanoSTING and varied the dose of treatment based on the duration of response that we have documented (**Figure 2**). The Alpha VOC (B.1.1.7) is known to be resistant to IFN-1 signaling *in vitro* and thus provides a challenging model to test the efficacy of NanoSTING^40, 41^. We pre-treated Syrian golden hamsters (n=6/group) with two intranasal doses of NanoSTING (30 μg and 120 μg) and 24 h later challenged the hamsters with ∼3 x 10^4^ CCID50 of the Alpha VOC (**Figure 6A**). Treatment with either dose of NanoSTING protected the hamsters from severe weight loss (**Figure 6B**). We used an integrated scoring rubric (range from 1-12) that accounts for histopathology of the lung tissue on day six after the viral challenge. We observed that NanoSTING treated hamsters had significant reductions in aggregate pathology score with minimal evidence of invasion by inflammatory cells or alveolar damage (**Figures 6C-D**). In addition, we quantified the viral titers in the lungs and nasal compartments and observed a significant reduction of viral titers in both compartments as early as day 2 post-challenge (**Figure 6E-F**). Thus, treatment with intranasal NanoSTING reduces *in vivo* replication of SARS-CoV-2 by orders of magnitude and confers protection against IFN-I evasive strains of SARS-CoV-2.

**Figure 6:**
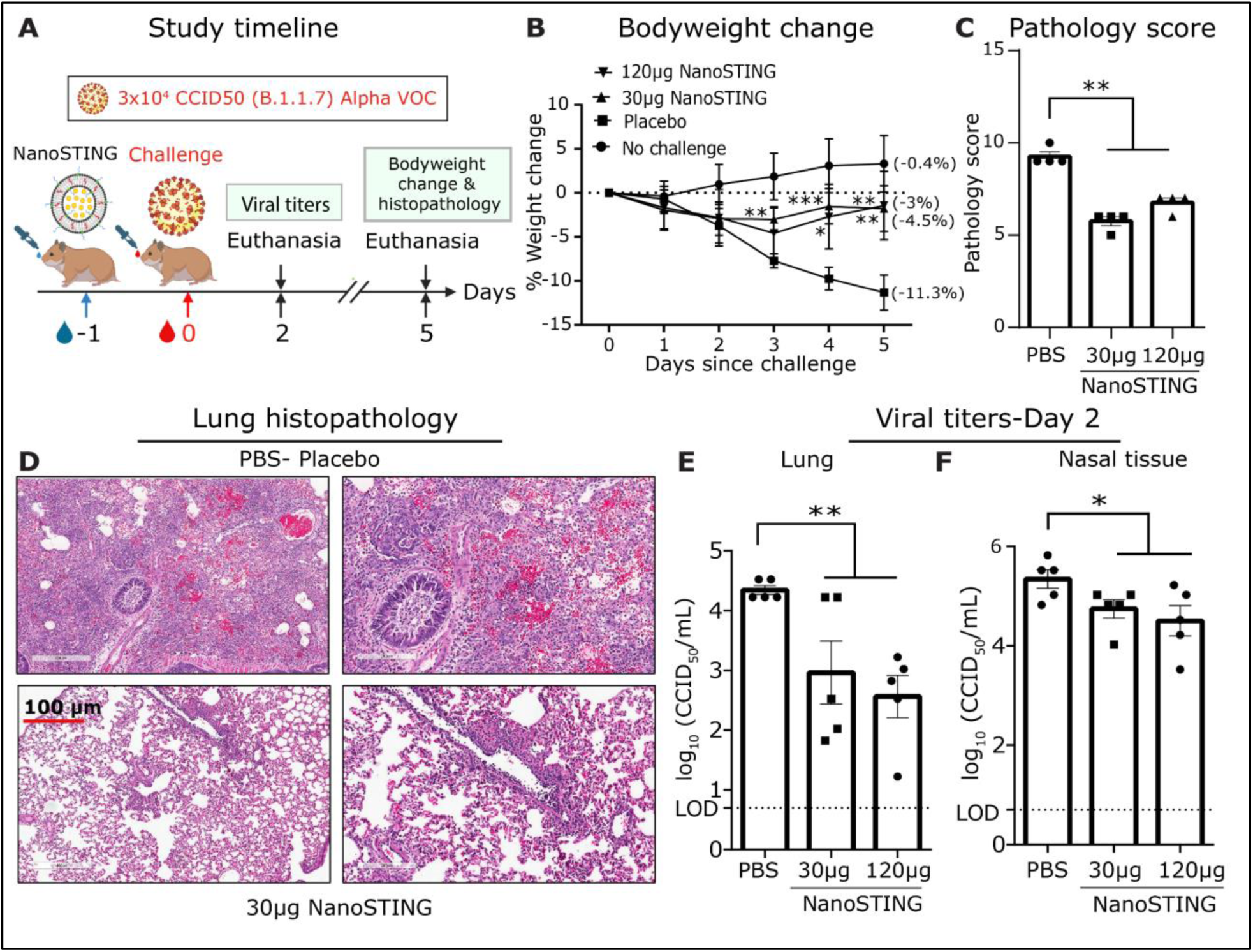
Protective efficacy of NanoSTING against the IFN evasive SARS-CoV-2 Alpha VOC (B.1.1.7) (A) We treated groups of 6 hamsters, each with two different doses of NanoSTING (30 µg and 120 µg) and 24 h later challenged with the ∼3 x 10^4^ CCID of SARS-CoV-2 Alpha VOC (B.1.1.7). We monitored animal weight changes daily for 5 days. Animals were euthanized for histopathology on day 5, with viral titers of lung and nasal tissues measured on day 2. (B) Change in bodyweight of hamsters. (C,D) Pathology score and a representative hematoxylin and eosin (H&E) image of the lung showing histopathological changes in lungs of hamsters treated with NanoSTING (30 µg) and PBS; all images were acquired at 20×; scale bar, 100 µm. (E, F) Viral titers were quantified in the lung and nasal tissue by plaque assay on day 2 after the challenge. The dotted line indicates limit of detection of the assay (LOD). *For viral titers and lung histopathology data, analysis was performed using a Mann-Whitney test. Vertical bars show mean values with error bar representing SEM. Each dot represents an individual hamster. Weight data was compared via mixed-effects model for repeated measures analysis. Lines depict group mean body weight change from day 0; error bars represent SEM. Asterisks indicate significance compared to the placebo-treated animals at each time point.* *\*\*\*\*p < 0.0001; ***p < 0.001; **p < 0.01; *p < 0.05; ns: not significant*.

### NanoSTING treatment prevents infection in hamsters exposed to the SARS-CoV-2 Omicron VOC

The Omicron VOC (B.1.1.529) is among the most infectious strains of SARS-CoV-2. Using the Omicron VOC sets up the highest bar for NanoSTING to prevent viral spread. We set up a transmission experiment designed to answer two fundamental questions: (1) does the prophylactic treatment of infected (index) hamsters prevent transmission to contact hamsters, and (2) does the post-exposure treatment of contact hamsters mitigate viral replication? Accordingly, we set up an experiment with three groups (n=16) of hamsters. In each group, eight index hamsters were intranasally infected with the ∼3 x 10^4^ CCID50 of the SARS-CoV-2 Omicron VOC, and one day after infection, each index hamster was paired with a cohoused contact hamster (**Figure 7A**). Consistent with the known milder disease of the Omicron VOC, none of the infected animals showed weight loss (**Figure S9**)^42^. We quantified the viral titers in the contact hamsters that were: (a) cohoused with placebo-treated infected index hamsters (group 1), (b) cohoused with NanoSTING (120 µg) treated index hamsters (group 2), or (c) treated with NanoSTING after cohousing with infected but untreated hamsters (group 3). As with the other strains of SARS-CoV-2 that we tested, NanoSTING pre-treatment of the infected hamsters almost completely blocked transmission (7/8 animals treated were virus-free vs 1/8 untreated animals were virus-free). Significantly, post-exposure treatment of the contact hamsters was also extremely effective at preventing infection (6/8 animals treated were virus-free), and all animals demonstrated a significant reduction in viral titers (**Figures 7B-C)**. These results directly demonstrate that NanoSTING is highly effective at blocking transmission even with the highly infectious Omicron VOC.

**Figure 7:**
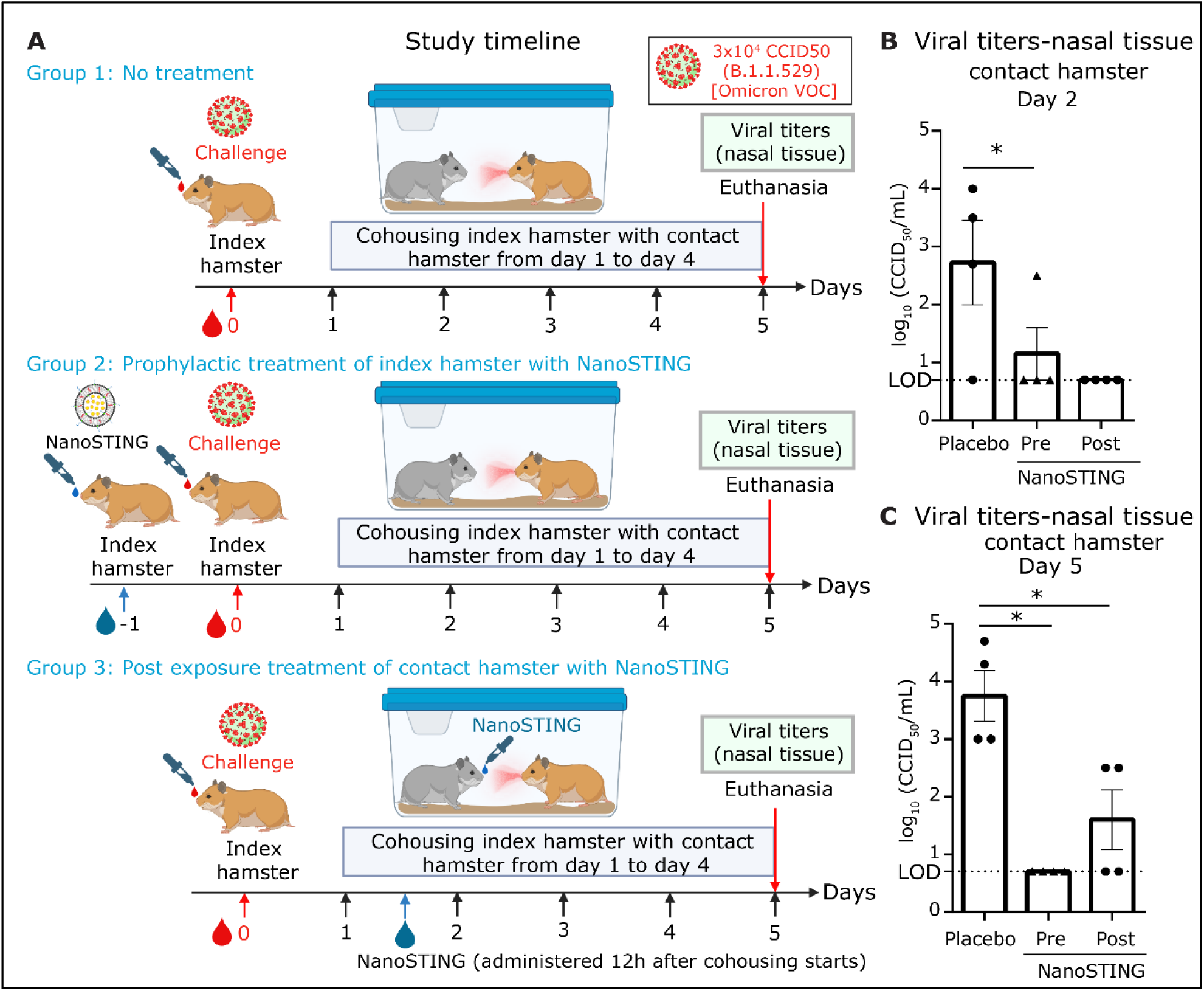
Intranasal administration of NanoSTING limits transmission and viral replication in the nasal passage of contact hamsters exposed to the SARS-CoV-2 Omicron (B.1.1.529) VOC. (A) Experimental set up: For group1, we challenged groups of 8 hamsters each on day 0 with ∼3 x 10^4^ of SARS-CoV-2 Omicron VOC (B.1.1.529) and after 24 h cohoused index hamsters in pairs with contact hamsters (n=8) for 4 days in clean cages. In group 2, we pre-treated the hamsters with 120 µg of NanoSTING 24 h prior to infection, and in group 3, we treated the contact hamsters with NanoSTING 12 h after the cohousing period began. We euthanized the contact and index hamsters on day 4 of cohousing. Viral titers in the nasal tissue of the index and contact hamsters were used as primary endpoints. (B, C) Infectious viral particles in the nasal tissue of contact hamsters at day 2 and day 5 after viral administration post-infection were measured by plaque assay. The dotted line indicates limit of detection of the assay (LOD). See also Figure S9 *For viral titers, analysis was performed using a Mann-Whitney test. Vertical bars show mean values with error bar representing SEM. Each dot represents an individual hamster. Mann-Whitney test: ****p < 0.0001; ***p < 0.001; **p < 0.01; *p < 0.05; ns: not significant*.

### Treatment with NanoSTING induces protection from influenza superior to oseltamivir

Influenza viruses have evolved multiple mechanisms to dampen the host’s innate immunity, including the attenuation of interferon responses by the NS1 protein^43, 44^. One of the primary treatment options against influenza involves post-exposure prophylaxis using oseltamivir which inhibits the influenza neuraminidase protein. We thus compared the efficacy of NanoSTING in comparison to oseltamivir in mouse models of influenza.

We challenged groups of ten mice with 2 x 10^4^ CCID of Influenza A/California/04/2009 (H1N1dpm). We treated them with a clinically relevant dose of oseltamivir (30 mg/kg/day), twice daily, for five days (**Figure S10A**)^45^. The untreated animals started losing significant weight by day three, and showed a mean peak weight loss of 31.1 % (**Figure S10B**). By contrast, animals treated with oseltamivir were moderately protected, showing a mean peak weight loss of 21.3% (**Figure S10B**). We next compared prophylactic treatment with either oseltamivir (two doses of 30mg/kg/day) or NanoSTING (single dose at 40 µg) followed by challenge with 2 x 10^4^ CCID of H1N1dpm (**Figure 8A**). Prophylactic administration of oseltamivir was ineffective, as animals in the placebo (mean peak weight loss of 27.6 %) and oseltamivir treated (mean peak weight loss 32.6 %) groups showed marked weight loss (**Figure 8B**). By comparison, a single dose of NanoSTING offered strong longitudinal protection from weight loss (mean peak weight loss 14.7 %). These results demonstrate that prophylactic treatment with NanoSTING is superior to oseltamivir treatment.

**Figure 8:**
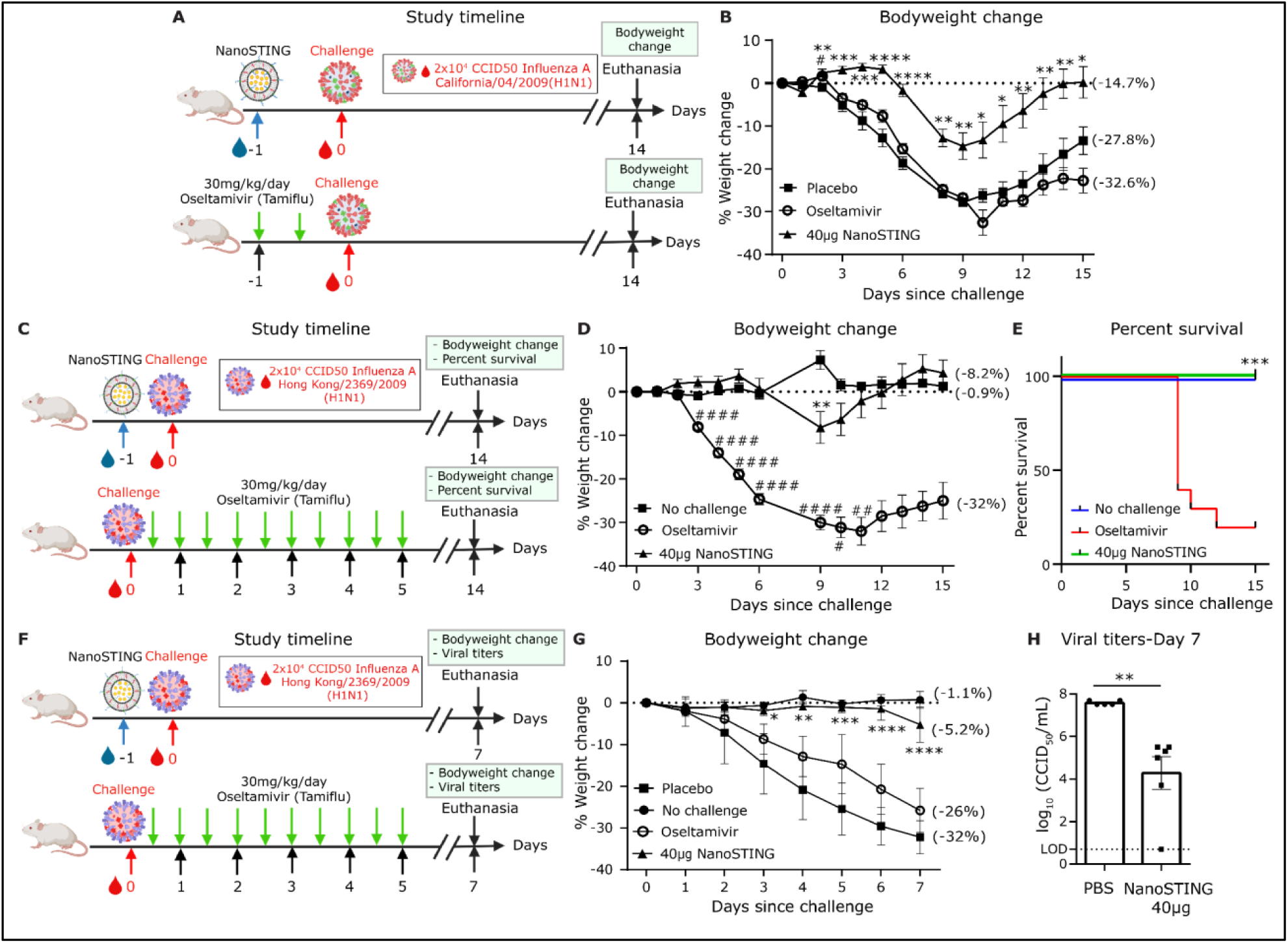
NanoSTING offers protection against Oseltamivir-sensitive and resistant strains of Influenza A. (A) Experimental set up: We treated groups of 10 BALB/c mice, each with a single dose of NanoSTING (40 µg) and Oseltamivir (30mg/kg/day-administered twice daily) or placebo and 24 h later challenged with 2 x 10^4^ CCID of Influenza A/California/04/2009 (H1N1dpm) strain and monitored for 14 days. Bodyweight change was used as the primary end point. Oseltamivir was used as a control. (B) Percent bodyweight change for the different groups of mice. (C) Experimental set up: We treated groups of 10 BALB/c mice with a single intranasal dose of NanoSTING (40 µg) and 24h later challenged with 2 x 10^4^ CCID of influenza A/Hong Kong/2369/2009 (H1N1)-H275Y [A-H275Y]. We evaluated the animals for 14 days and used weight loss and survival as the primary endpoints. We treated one group of mice with a clinically relevant dose of oseltamivir, twice daily, for five days. (D) Percent weight change compared to the weight at d0 at the indicated time intervals post-infection. (E) Survival of the different groups of mice. (F) Experimental set up: We treated groups of 10 BALB/c mice with a single intranasal dose of NanoSTING (40 µg), and 24h later challenged with 2 x 10^4^ CCID of influenza A/Hong Kong/2369/2009 (H1N1)-H275Y [A-H275Y]. We monitored the animals for 7 days for body weight change and quantified viral titers at the end of the study. We treated one group of mice with oseltamivir, twice daily, for five days. (G) Weight change of the different groups of mice. (H) Viral titers were measured by plaque assay in lungs post 7 days after infection. The dotted line indicates limit of detection of the assay (LOD). *For viral titers, analysis was performed using a Mann-Whitney test. Vertical bars show mean values with error bar representing SEM. Each dot represents an individual mouse. Weight data was compared via mixed-effects model for repeated measures analysis. Lines depict group mean body weight change from day 0; error bars represent SEM. For Figures 8B and 8G, asterisks indicate statistical significant differences between NanoSTING-treated group and placebo-treated animals, whereas, pound sign show statistical significant differences between Oseltamivir-treated group and placebo-treated animals. For Figure 8D, asterisks indicate statistical significant differences between NanoSTING-treated group and non-challenged animals, whereas, pound sign indicate statistical significant differences between Oseltamivir-treated group and non-challenged animals. We compared survival percentages between NanoSTING-treated and Oseltamivir-treated animals using the Log-Rank Test (Mantel-Cox)*. *\*\*\*\*p < 0.0001; ***p < 0.001; **p < 0.01; *p < 0.05; ns: not significant*.

The evolution of resistance to treatment is predictable and common with influenza. A single amino acid mutation (His275Tyr) with neuraminidase has led to oseltamivir-resistant influenza viruses in humans^46^. Since NanoSTING relies on the host’s innate immune response and should be effective against treatment resistant strains, we next evaluated its potency against oseltamivir-resistant influenza A in mice. We treated groups of ten mice with a single intranasal dose of NanoSTING (40 µg) and 24 h later challenged with 2 x 10^4^ CCID of influenza A/Hong Kong/2369/2009 (H1N1)-H275Y [A-H275Y] (**Figure 8C**). We evaluated the animals for 14 days and used weight loss and survival as the primary endpoints. We treated one group of mice with oseltamivir (30 mg/kg/day), twice daily, for five days, as a control^45^. NanoSTING treated animals were well-protected from weight loss (mean peak weight loss of 8.2%) in comparison to oseltamivir treatment (mean peak weight loss of 32%) [**Figure 8D**]. The weight loss in the NanoSTING treated animals was transient between days 6-10, and outside of this window, the weight loss in the animals was no different from unchallenged animals (**Figure 8D**). By contrast, starting at day 4, oseltamivir-treated animals showed significant weight loss until the end of the study (day 15). Consistent with these observations, 100% of NanoSTING treated animals survived, whereas only 20% of oseltamivir treated animals survived (**Figure 8E**). Collectively these results further reinforce the efficacy of NanoSTING treatment against multiple strains of influenza.

To test the impact of NanoSTING treatment on viral titers within the lung, we repeated the challenge experiments with influenza A-H275Y and euthanized the animals at day 7 (**Figure 8F**). A single-dose treatment with NanoSTING again protected animals from weight loss (mean peak weight loss at day 7 of 5.2 % vs. 32% placebo) [**Figure 8G**]. In addition, infectious particles in the lung 7 days after viral exposure were reduced by 500-fold compared to the placebo-treated group accounting for the ability of NanoSTING to help prevent disease and death (**Figure 8H**). In aggregate, these experiments confirmed that NanoSTING works as a broad-spectrum antiviral against influenza by protecting from weight loss, reducing viral titers, and preventing death.

## Discussion

The availability of prophylactics or early post-exposure therapeutics (treatment initiated prior to clinical symptoms) that can both prevent disease and reduce transmission is an urgent and unmet clinical need. Here, we have demonstrated that a single dose of intranasal NanoSTING can work as prophylactic and therapeutic against multiple respiratory viruses (and standard treatment-resistant variants).

During the last two years, SARS-CoV-2 has spread at an alarming rate, and the U.S. in on track to record one million confirmed deaths from COVID in a few weeks. The current pandemic has once again highlighted that our therapeutic arsenal against RNA viruses is inadequate. Vaccines are our preferred means of protection against SARS-CoV-2, but they suffer from three drawbacks. First, while the current generation of vaccines were tested at remarkable speed, even this rate of development lags as vaccines need to be custom manufactured for each emerging virus. Second, the mutational plasticity of RNA viruses like SARS-CoV-2 facilitates their evolution, and newer variants with immune escape potential have emerged. This necessitates ongoing booster shots in adults to achieve at least transitory complete protection from disease, even as the entire human population is not yet fully vaccinated from SARS-CoV-2^47, 48^. As the human experience with influenza has illustrated, requiring additional booster shots reduces human compliance that in turn facilitates the spread of disease. Compounding this problem is that immunosuppressed vaccine recipients fail to be sufficiently protected, and reservoirs are emerging for SARS-CoV-2 outside of humans^49^. Third, despite the efficacy of the current intramuscular vaccines in preventing disease, they do not prevent transmission^50^. The evolution of the SARS-CoV-2 Omicron (B.1.1.529) VOC shows that viruses can quickly adapt to facilitate rapid spread using the nasal cavity as a sanctuary. Thus, while vaccines are necessary, they are not sufficient to fight RNA viruses.

Monoclonal antibodies targeting viruses, like vaccines, offer protection against respiratory disease, but suffer from the same disadvantages of vaccines listed above. Furthermore, their window of use is limited as the emergence of SARS-CoV-2 VOCs such as Omicron can quickly render them ineffective^47^. Additionally, monoclonal antibodies are expensive and administered in a clinical setting, further limiting their widespread use. NanoSTING offers an alternative by generating an immune response which appears advantageous to instilling immunity. Focusing on efficacy, intravenous prophylactic administration of antibody (12 h before challenge) in hamsters led to protection from clinical disease (∼2-5 % weight loss and ∼300-fold reduction in viral titers in the lung) albeit with no impact on transmission^51, 52^. NanoSTING (which is easier to administer) provides a broader window of administration (24-72 h), with comparable efficacy in reducing clinical disease while also reducing transmission.

Oral antivirals that directly inhibit one or more viral proteins have been developed against SARS-CoV-2 (*e.g.* paxlovid and molnupiravir) and Influenza (Oseltamivir) are approved for use in humans, but are also susceptible to viral evolution and resistance^8^. Furthermore, these therapeutics are designed as oral post-exposure prophylactics to prevent clinical disease and have no impact on viral transmission^53^. In contrast to these pathogen-specific drugs, NanoSTING works broadly against multiple respiratory viruses, including oseltamivir-resistant influenza highlighting its translational potential. Furthermore, the superior efficacy of NanoSTING compared to paxlovid and molnupiravir in small animal models and the fact that these antivirals are efficacious in humans (30-89% in reducing clinical disease with SARS-CoV-2) augers well for the clinical potential of NanoSTING^54^.

Immunomodulators, including defective viral genome particles, cytokines, and small molecule agonists, have been tested as antivirals. Defective interfering particles (DIPs) have incomplete genomes and, when administered therapeutically, inhibit replication of the wild-type virus^55^. Although these particles have demonstrated efficacy for SARS-CoV-2 and Influenza in mitigating disease in small animal models, the DIPs must be generated for each virus individually^55, 56^. Defective viral genomes (DVGs) based on the poliovirus induce a broad IFN-I response and are protective against multiple viruses^57^. However, DVGs need to replicate *in vivo* after administration, and this limited replication is essential for their efficacy. However, their broad applicability is limited by concerns about both safety and the presence of pre-existing antibodies in vaccinated people. Lipid nanoparticles complexed with the defective genomes can mitigate these concerns and have shown efficacy against SARS-CoV-2 VOCs in K18-hACE2 mice; the generalizability of this approach in the absence of viral replication to other viruses has not been demonstrated^57^.

Direct administration of aerosolized interferons to engage antiviral innate immunity has been tested in animals and humans. In hamsters challenged with SARS-CoV-2, prophylactic or early administration of universal IFN reduces lung damage, provides moderate protection against weight loss (10% vs. 20% for untreated animals), and reduces infectious viral particles (100-fold)^58, 59^. NanoSTING appears to offer superior efficacy when compared to these data. In humans, post-exposure prophylaxis with nebulized IFN-α2b was associated with reduced in-hospital mortality compared to no administration of IFN-α2b. By contrast, administration of IFN-α2b more than five days after admission delayed recovery and increased mortality, suggesting that the timing of IFN-α2b administration is critical for efficacy ^60^. The limited impact of IFN-α for COVID-19 mirrors its negligible efficacy as a prophylactic against Influenza in humans^61^. Other synthetic small molecule agonists of pattern recognition receptors (PRRs), including stem-loop RNA 14 (SLR14), a minimal RIG-I (Retinoic acid-inducible gene I) agonist, and STING agonist, diAbzl, have been tested against SARS-CoV-2 in K18-hACE2 mice^22, 41^. As with all small-molecule drugs, their safety, off-target activity, and pharmacokinetics must be thoroughly evaluated before translation. NanoSTING is comprised of naturally occurring lipids that have already been tested in humans and cGAMP, the immunotransmitter of danger signals that are conserved across mammals, including humans^62^. As illustrated, NanoSTING leads to safe and sustained delivery and consequently functions as a broad-spectrum antiviral.

Our data illustrate that NanoSTING has emerged as a first-in-class immunoantiviral as it is safe, stable, and effective against multiple viruses and variants. It achieves its antiviral effects by rapidly engaging and sustaining activation of the STING pathway. Indeed, an advantage of using the natural immunotransmitter, cGAMP is that STING activation can lead to both IFN dependent and independent activities to control viral replication^19, 20, 62^. NanoSTING exhibits a broad spectrum of activity against existing viruses, but by activating the innate response, it protects against existing’s viruses and likely emerging threats. Furthermore, by enabling immunological memory, NanoSTING minimizes clinical symptoms during primary infection while preserving durable protection from reinfection without the need for retreatment. We envision NanoSTING as a treatment to prevent respiratory viral disease in vulnerable populations or to rapidly intervene in respiratory infections before etiology is determined.

## Materials & Methods

### Preparation of NanoSTING

The liposomes contained DPPC, DPPG, Cholesterol (Chol), and DPPE-PEG2000 (Avanti Polar lipids) in a molar ratio of 10:1:1:1. To prepare the liposomes, we mixed the lipids in CH_3_OH and CHCl_3_, and we evaporated them at 45°C using a vacuum rotary evaporator. The resulting lipid thin film was dried in a hood to remove any residual organic solvent. We hydrated the lipid film by adding a pre-warmed cGAMP (Chemietek) solution (3 mg/mL in PBS buffer at pH 7.4). We mixed the hydrated lipids at an elevated temperature of 65°C for an additional 30 min, and subjected them to freeze-thaw cycles. We next sonicated the mixture for 60 min using a Branson Sonicator (40 kHz) and used Amicon Ultrafiltration units (MW cut off 10 kDa) for removing the free untrapped cGAMP. Finally, we washed the NanoSTING (liposomally encapsulated STINGa) three times using PBS buffer. We measured the cGAMP concentration in the filtrates against a calibration curve of cGAMP at 260 nm using Take3 Micro-Volume absorbance analyzer of Cytation 5 (BioTek). We calculated the final concentration of cGAMP in NanoSTING and encapsulation efficiency by subtracting the concentration of free drug in the filtrate.

For checking the stability, we stored the NanoSTING at 24°C and 37°C for 1, 2, 3, 7, 14, and 30 days. We measured the average hydrodynamic diameter and zeta potential of liposomal particles using DLS and zeta sizer on Litesizer 500 (Anton Paar).

### Cell lines

THP-1 dual cell line (Invivogen) was cultured in a humidified incubator at 37°C and 5% CO_2_ and grown in RPMI/10% FBS (Corning, NY, USA). In addition, we supplemented the THP-1 dual cell line with the respective selection agents (100 μg/mL zeocin + 10 μg/mL blasticidin) and the corresponding selection cytostatics from Invivogen.

### Cell stimulation experiments with luciferase reporter enzyme detection

We performed the cell stimulation experiments using the manufacturer’s instructions (Invivogen, CA, USA). First, we seeded the cells in 96 well plate at 1 x 10^5^ cells/well in 180 μL growth medium. Next, we made serial dilutions of NanoSTING on a separate plate at concentrations ranging from 2.5-10 µg/mL in the growth medium. We then incubated the cells at 37°C for 24 h. For detecting IRF activity, we collected 10 μL of culture supernatant/well at time points of 6 h, 12 h, and 24 h and transferred it to a white (opaque) 96 well plate. Next, we read the plate on Cytation 7 (Cytation 7, Bio-Tek Instruments, Inc.) after adding 50 μL QUANTI-Luc™ (Invivogen) substrate solution per well, followed by immediate luminescence measurement. The data was recorded as relative light units (RLU).

### Mice and NanoSTING treatment

All studies using animal experiments were reviewed and approved by University of Houston (UH) IACUC. We purchased the female 7 to 9-week-old BALB/c mice from Charles River Laboratories. Gender/sex as a variable was not tested. We treated the groups of BALB/c mice (n=3-4) intranasally with varying amounts of NanoSTING (10 to 40 μg) after sedating them with isoflurane. We euthanized the animals after 6 h, 12 h, 24 h, 36 h, and 48 h and harvested blood, nasal turbinates, and lungs. We kept the blood at room temperature (RT) for 10 min to facilitate clotting and centrifuged it for 5 min at 2000g. We collected the serum, stored it at −80°C, and used it for ELISA.

### ELISA

We homogenized nasal turbinates and lung tissue samples in 1:20 (w/v) of tissue protein extraction reagent (Thermo Fisher, # 78510), then centrifuged them to pellet tissue debris. We centrifuged the homogenates for 10 min at 2500g. We assayed the supernatants for cGAMP, IFN-β, and CXCL10 using quantitative ELISA. cGAMP ELISA was performed using 2’3’-cGAMP ELISA kit (Cayman Chemicals, MI, USA). IFN-β concentrations were tested using mouse IFN-beta Quantikine ELISA kit (R&D Systems, MN, USA). Mouse IP-10 ELISA kit (CXCL10) was used to perform the CXCL10 ELISAs (Abcam, MA, USA). cGAMP, IFN-β, and CXCL10 concentrations were tested by titering 30 µg total protein from nasal turbinates and lung lysates. All serum samples were tested at 50X dilutions to test cGAMP, IFN-β, and CXCL10 concentrations.

### RNA isolation, cDNA preparation, and qRT-PCR

We excised mouse nasal turbinates tissues and placed approximately 20 mg of tissue in 2 mL tubes containing 500 μL RNeasy lysis buffer (RLT) and a single stainless steel bead. Next, we homogenized the tissue using a tissue lyser (Qiagen, Hilden, Germany) before total RNA extraction using an RNeasy kit (Qiagen, #74104), following the manufacturer’s instructions. Extracted RNA was DNase treated using a DNA-free DNA removal kit (Invitrogen, #AM1906). Next, 1 µg of total RNA was converted to cDNA using a High-Capacity cDNA reverse transcription kit (Invitrogen, #4368813). We diluted the resultant cDNA to 1:10 before analyzing quantitative real-time polymerase chain reaction (qRT-PCR). We performed qRT-PCR reaction using SsoFastTM EvaGreen® Supermix with Low ROX (Biorad, # 1725211) on AriaMx Real-time PCR System (Agilent Technologies, Santa Clara, CA). We normalized the results to GAPDH (glyceraldehyde-3-phosphate dehydrogenase). We determined the fold change using the 2-DDCt method, comparing treated mice to naive controls. See Table S2 for the list of primers sequences used in this study.

### Syrian Golden Hamsters

All studies using animal experiments were reviewed and approved by UH IACUC. We purchased the 6 to 10 weeks old male and female hamsters *(Mesocricetus auratus*) from Charles River Laboratories.

### Safety studies of NanoSTING on hamsters

We designed a pilot study to test if repeated administration of NanoSTING causes clinical symptoms (fever or weight loss). We administered group (n=4/group) of animals with daily doses of 60 µg of NanoSTING intranasally for four consecutive days. We used naive hamsters as controls (n=4/group). The animals were monitored daily for bodyweight change and body temperature. We euthanized the animals 24 h after administering the last dose and harvested lungs.

### Processing of the hamster’s lungs for qRT-PCR and mRNA sequencing

For isolation of single-cell suspension from the lungs, each lung was cut into 100–300 mm^2^ pieces using a scalpel. We transferred the minced tissue to a tube containing 5 mL of digestion Buffer containing collagenase D (2mg/mL, Roche #11088858001) and DNase (0.125 mg/mL, Sigma #DN25) in 5 mL of RPMI (Corning, NY, USA) for 1 h and 30 min at 37 °C in the water bath with vortexing after every 10 min. We disrupted the remaining intact tissue by passaging (6-8 times) through a 21-gauge needle. Post 1 h and 30 min of incubation, we added 500 µL of iced-stopping buffer (1x PBS, 0.1M EDTA) to each falcon tube to stop the reaction. We then removed tissue fragments and the majority of the dead cells with a 40 µm disposable cell strainer (Falcon, #352340), and we collected the cells after centrifugation. We lysed the red blood cells by resuspending the cell pellet in 3 mL of ACK lysing Buffer (Gibco, #A1049201) and incubated for 3 min at RT, followed by centrifugation. We discarded the supernatants and resuspended the cell pellets in 5 mL of complete RPMI medium (Corning, NY, USA). We enumerated lung cells by trypan blue exclusion.

### qRT-PCR and mRNA sequencing for hamster’s lung cells

Total RNA was extracted from whole lung cells using RNeasy kit (Qiagen, #74104), following the manufacturer’s instructions. Extracted RNA was DNase treated using a DNA-free DNA removal kit (Invitrogen, #AM1906). 1 µg of total RNA was converted to cDNA using a High-capacity cDNA reverse transcription kit (Invitrogen, #4368813). We diluted the resultant cDNA to 1:10 before analyzing qRT-PCR. We performed qRT-PCR reaction using SsoFastTM EvaGreen® Supermix with Low ROX (Biorad, #1725211) on AriaMx Real-time PCR System (Agilent Technologies, Santa Clara, CA). We normalized the results to *Actb* (β-actin gene). We determined the fold change using the 2-DDCt method, comparing treated mice to naive controls. The primer sequences are provided in Table S3. Preparation of RNA library and mRNA sequencing was conducted by Novogene Co., LTD (Beijing, China). We paired and trimmed the fastq files using Trimmomatic (v 0.39) and aligned them to the Syrian golden hamster genome (MesAur 1.0, ensembl) using STAR (v 2.7.9a). We determined the differential gene expression using DESeq2 (v 1.28.1) package^63^. To perform gene set enrichment analysis, we used a pre-ranked gene list of differentially expressed genes in GSEA software (UC San Diego and Broad Institute). To generate the gene set for IFN-independent activities of STING, we collected genes with a 2-fold change increase in BMDM-STING S365A-DMXAA vs BMDM-STING S365A-DMSO samples from the GSE149744 dataset as described previously^19^.

### Viruses and biosafety

*Viruses.* Isolates of SARS-CoV-2 were obtained from BEI Resources (Manassas, VA) and amplified in Vero E6 cells to create working stocks of the virus. Influenza A/California/04/2009 was kindly provided by Elena Govorkova (St. Jude Children’s Research Hospital, Memphis, TN) and was adapted to mice by Natalia Ilyushina and colleagues at the same institution. Influenza A/Hong Kong/2369/2009 (H1N1pdm) was provided by Kwok-Yung Yuen from The University of Hong Kong, Hong Kong Special Administrative Region, People’s Republic of China. The virus was adapted to mice by four serial passages in the lungs of mice and plaque purified at USU.

*Biosafety.* Studies with influenza virus were completed within the ABSL-2 space of the Laboratory Animal Research Center (LARC) at USU. Studies involving SARS-CoV-2 were completed within the ABSL-3 space of the LARC at USU.

### Viral challenge studies in animals

#### Animals

For SARS-Cov-2 animal studies completed at USU, 6 to 10-week-old male and female golden Syrian hamsters were purchased from Charles River Laboratories and housed in the ABSL-3 animal space within the LARC. For influenza virus animal studies, 8-week-old BALB/c mice were purchased from Charles River Laboratories.

#### Infection of animals

Hamsters were anesthetized with isoflurane and infected by intranasal instillation of 1 x 10^4.5^ CCID_50_ of SARS-CoV-2 in a 100 µl volume. Mice were also anesthetized with isoflurane and infected with a 1 x 10^4.3^ CCID_50_ dose of influenza virus in a 90 µl volume.

Titration of tissue samples. Lung tissue and nasal tissue samples from hamsters and lung tissue samples from mice were homogenized using a bead-mill homogenizer using in minimum essential media. Homogenized tissue samples were serially diluted in test medium and the virus quantified using an endpoint dilution assay on Vero E6 cells for SARS-CoV-2 and on MDCK cells for influenza virus. A 50% cell culture infectious dose was determined using the Reed-Muench equation^64^.

#### Ethics

The animal experiments at USU were conducted in accordance with an approved protocol by the Institutional Animal Care and Use Committee of Utah State University. The work was performed in the AAALAC-accredited LARC of the university in accordance with the National Institutes of Health Guide for the Care and Use of Laboratory Animals (8^th^ edition; 2011).

### Histopathology

Lungs of the Syrian golden hamsters were fixed in 10% neural buffered formalin overnight and then processed for paraffin embedding. The 4-μm sections were stained with hematoxylin and eosin for histopathological examinations. Images were scanned using an Aperio ImageScope.

### Quantification and statistical analysis

Statistical significance was assigned when P values were <0.05 using GraphPad Prism (v6.07). Tests, number of animals (n), mean values, and statistical comparison groups are indicated in the Figure legends. Analysis of fold changes in gene expression, ELISA, and differences in viral titers after NanoSTING treatment were determined using a Mann-Whitney test. Survival percentages were tested using the Log-Rank Test (Mantel-Cox). Bodyweight data at each time point was compared *via* the mixed-effects model for repeated measures analysis.

### Quantitative modeling

To quantify the kinetics of SARS-CoV-2 infection in the upper respiratory tract (URT) in the presence of NanoSTING, we modified the innate immune model described by Ke et al. ^34^. We added an additional coefficient to the term responsible for refractory responses in the set of governing ordinary differential equations (ODEs), as shown in Figure 4 and supplementary data (Figure S6). The mean population parameter values and initial values were taken from Ke et al. ^34^. We solved the system of ODEs for different efficacies, treatment initiation time, and duration of response of NanoSTING using the ODE45 function in MATLAB 2018b. A sample MATLAB code for solving the system of equations has been provided in supplementary methods.

## Acknowledgments

This publication was supported by the NIH (R01GM143243), Owens foundation, and AuraVax Therapeutics. Parts of the figures were created using BioRender (<u>BioRender.com</u>).

## Author contributions

NV conceived the study. NV, AL, LC, MS, and BH designed the study. AL, AS, SRS, MK, MM, RK, SB, BH, and XL performed experiments. AL, AS, SRS, MK, BH, XL, and NV analyzed the data. MK performed modeling, and MF performed bioinformatic analyses. NV and AL drafted the manuscript and all authors contributed to the review and editing of the manuscript.

## Supplementary Figures

**Figure S1:**
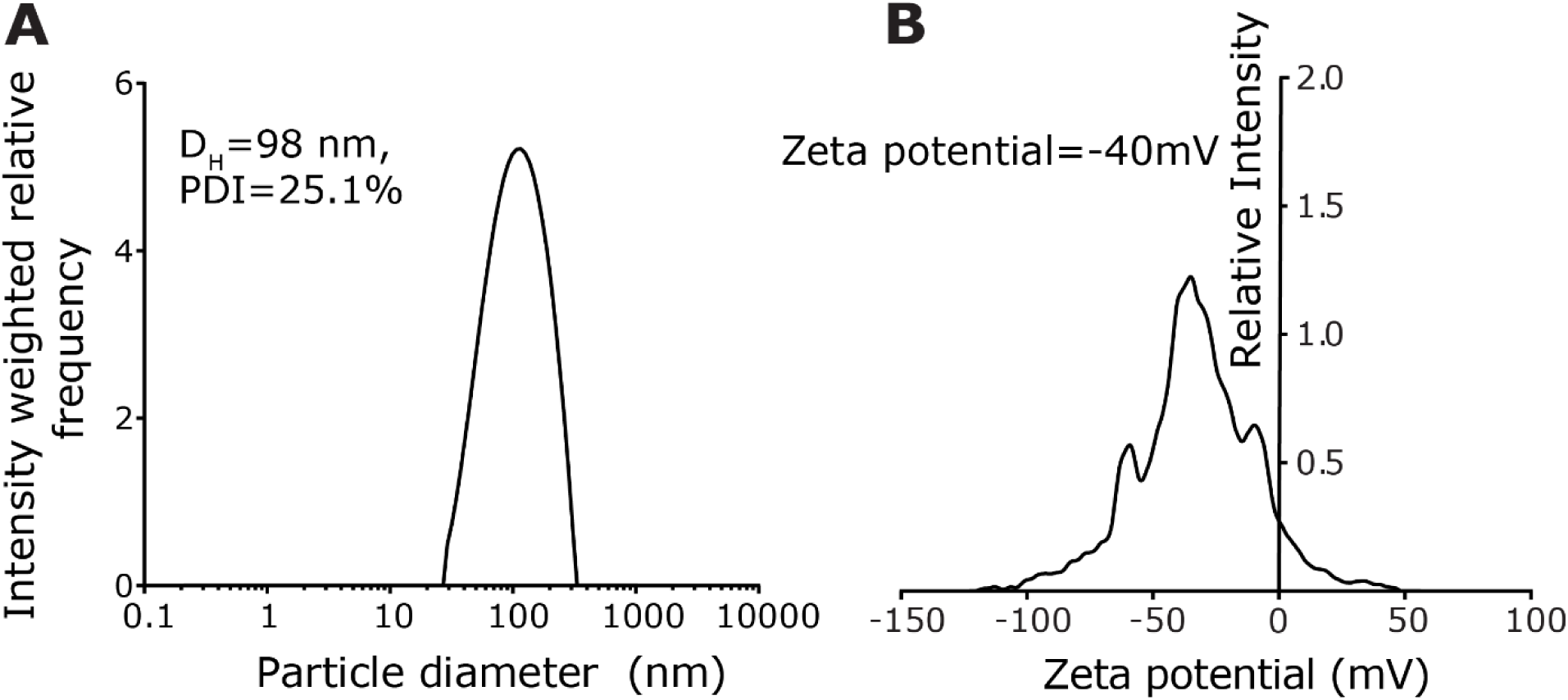
Characterization of NanoSTING (related to Figure 1) (A) Size distribution of NanoSTING measured by DLS. (B) Zeta potential of the NanoSTING measured by electrophoretic light scattering (ELS).

**Figure S2:**
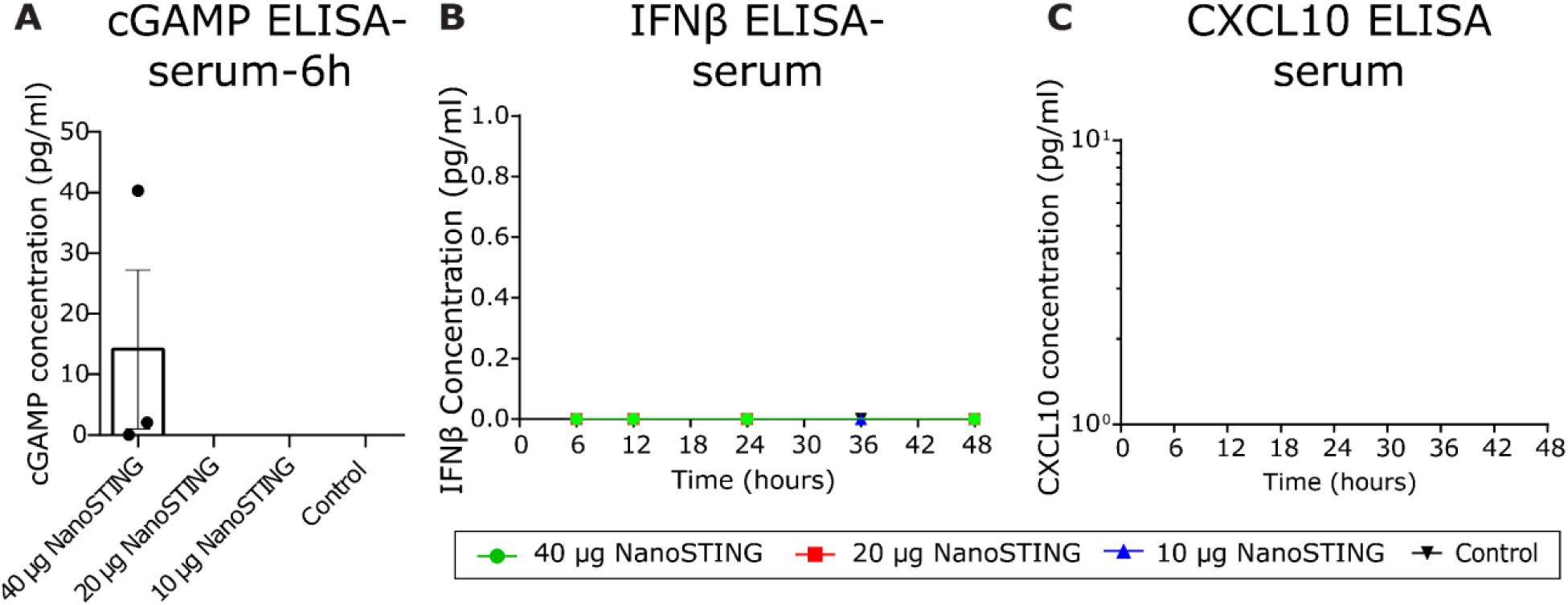
ELISA results confirmed that stimulation of the innate immunity was localized and not systemic (related to **Figure 2**) (A) Quantification of cGAMP in the mouse serum after treatment with NanoSTING. (B) Detection of IFN-β concentration in the mouse serum using quantitative ELISA. (C) Detection of CXCL10 levels in the mouse serum using quantitative ELISA. *The analysis was performed using a Mann-Whitney test. Vertical bars show mean values with error bar representing SEM. Mann-Whitney test: ****p < 0.0001; ***p < 0.001; **p < 0.01; *p < 0.05; ns, not significant*.

**Figure S3:**
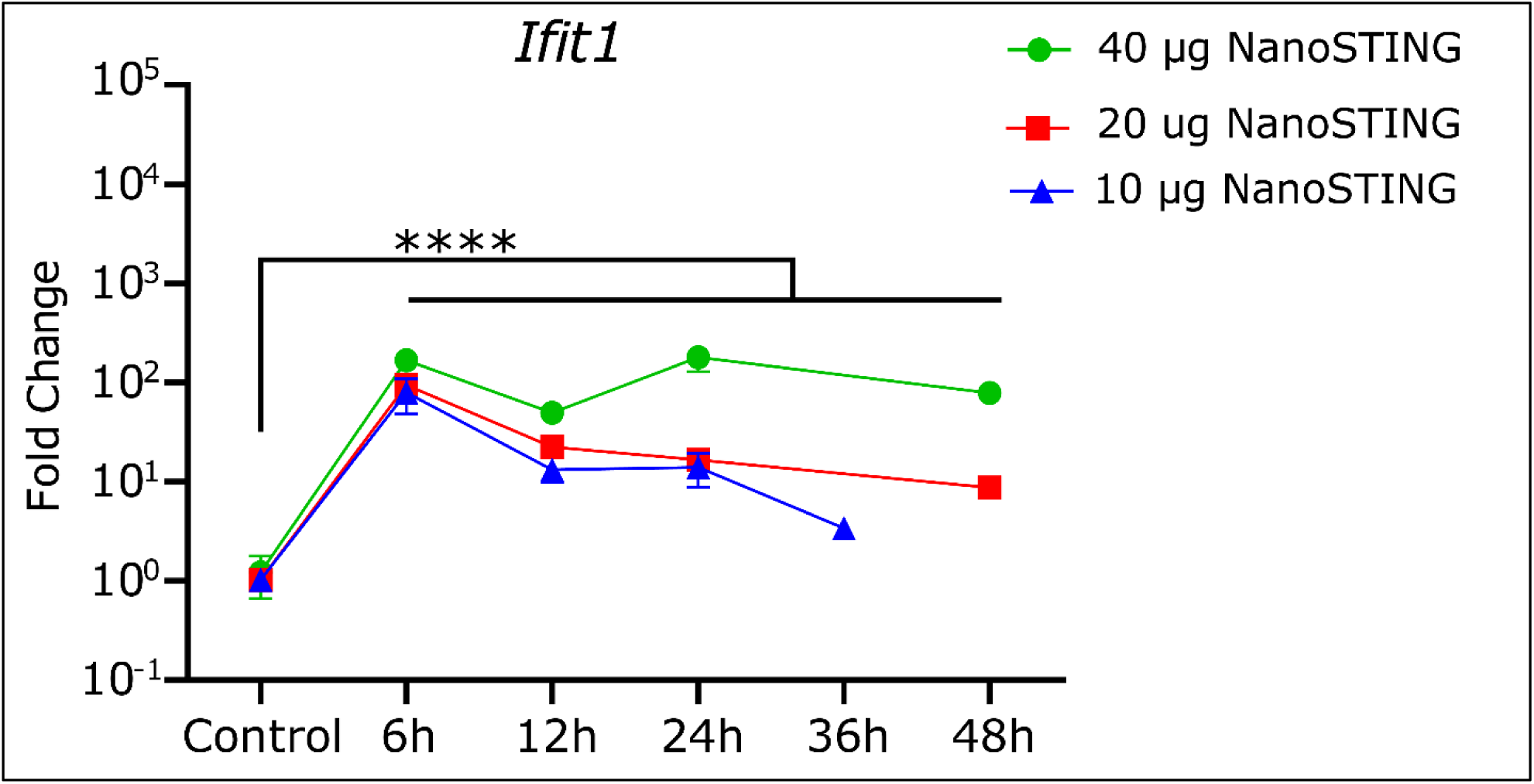
Real-time qRT-PCR for fold induction of *Ifit1* mRNA from NanoSTING treated mice compared with control mice nasal turbinates (related to **Figure 2**) *For fold changes in gene expression, analysis was performed using a Mann-Whitney test. Mann-Whitney test: ****p < 0.0001; ***p < 0.001; **p < 0.01; *p < 0.05; ns: not significant*

**Figure S4:**
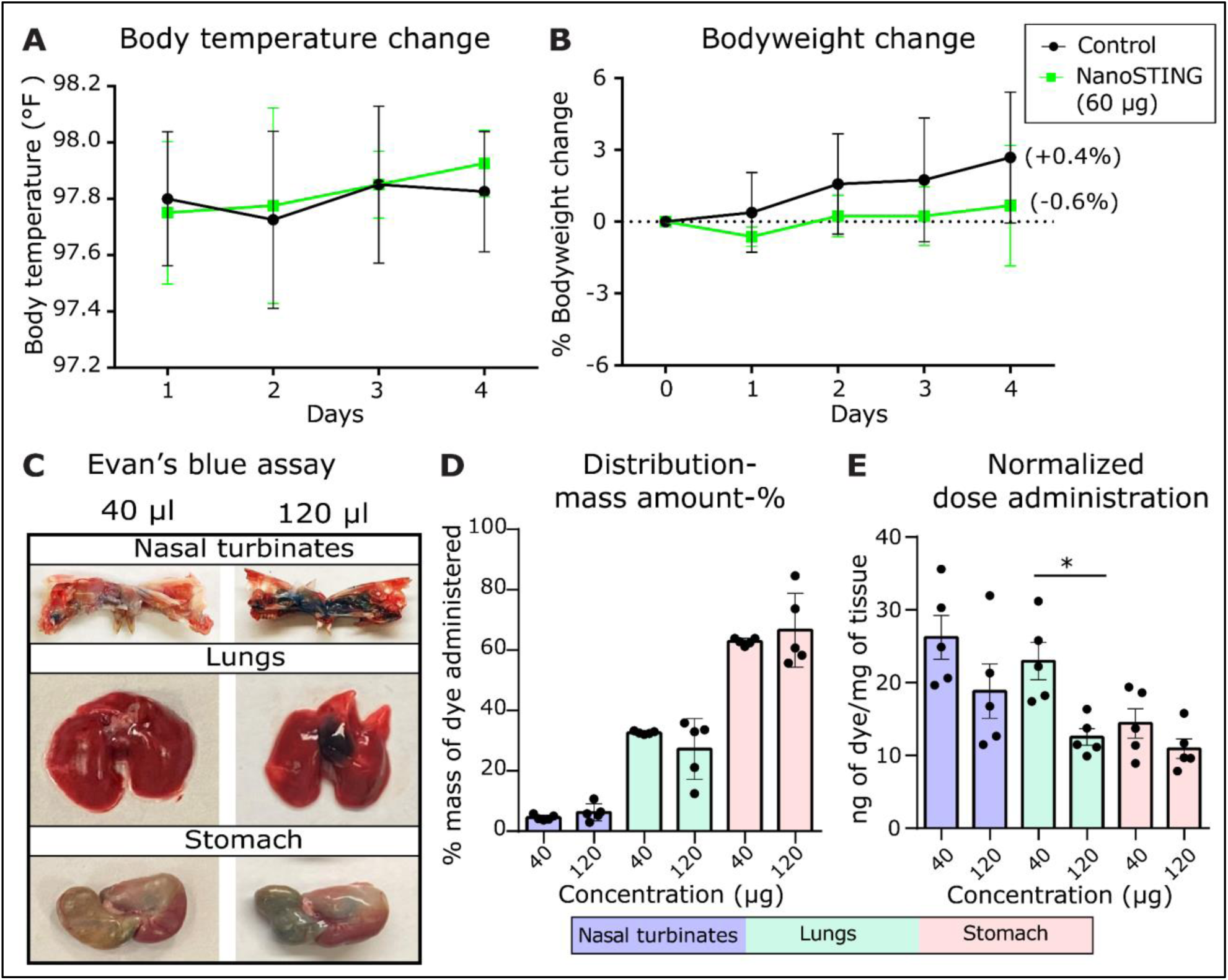
Safety and distribution studies for NanoSTING (related to **Figure 3**) (A, B) Safety studies of NanoSTING on hamster model. We administered groups of animals with daily doses of 60 µg of NanoSTING intranasally (n=4/group) or PBS (n=4/group) for four consecutive days. We monitored the hamsters daily for bodyweight change and body temperature change. We euthanized the hamsters on Day 5 after administering the last dose on Day 4, followed by the collection of lungs. Bodyweight change, body temperature change, qRT-PCR for lungs (Figure S5), and mRNA sequencing (Figure 3) were primary endpoints. (C) Evan’s blue dye assay. Hamsters (n=4/group) were intranasally administered with 0.125% Evans blue dye in PBS (40 μL and 120 μL). Representative images of nasal turbinates, lungs, and stomach dissected 2 min later are shown. (D, E) Distribution of Evan’s blue dye after intranasal administration. We treated the supernatants from homogenized lungs and stomach with trichloroacetic acid and analyzed absorbance at 620 nm. We interpolated the concentrations of dye from a standard curve. *The analysis was performed using a Mann-Whitney test. Vertical bars show mean values with error bar representing SEM. Mann-Whitney test: ****p < 0.0001; ***p < 0.001; **p < 0.01; *p < 0.05; ns: not significant*

**Figure S5:**
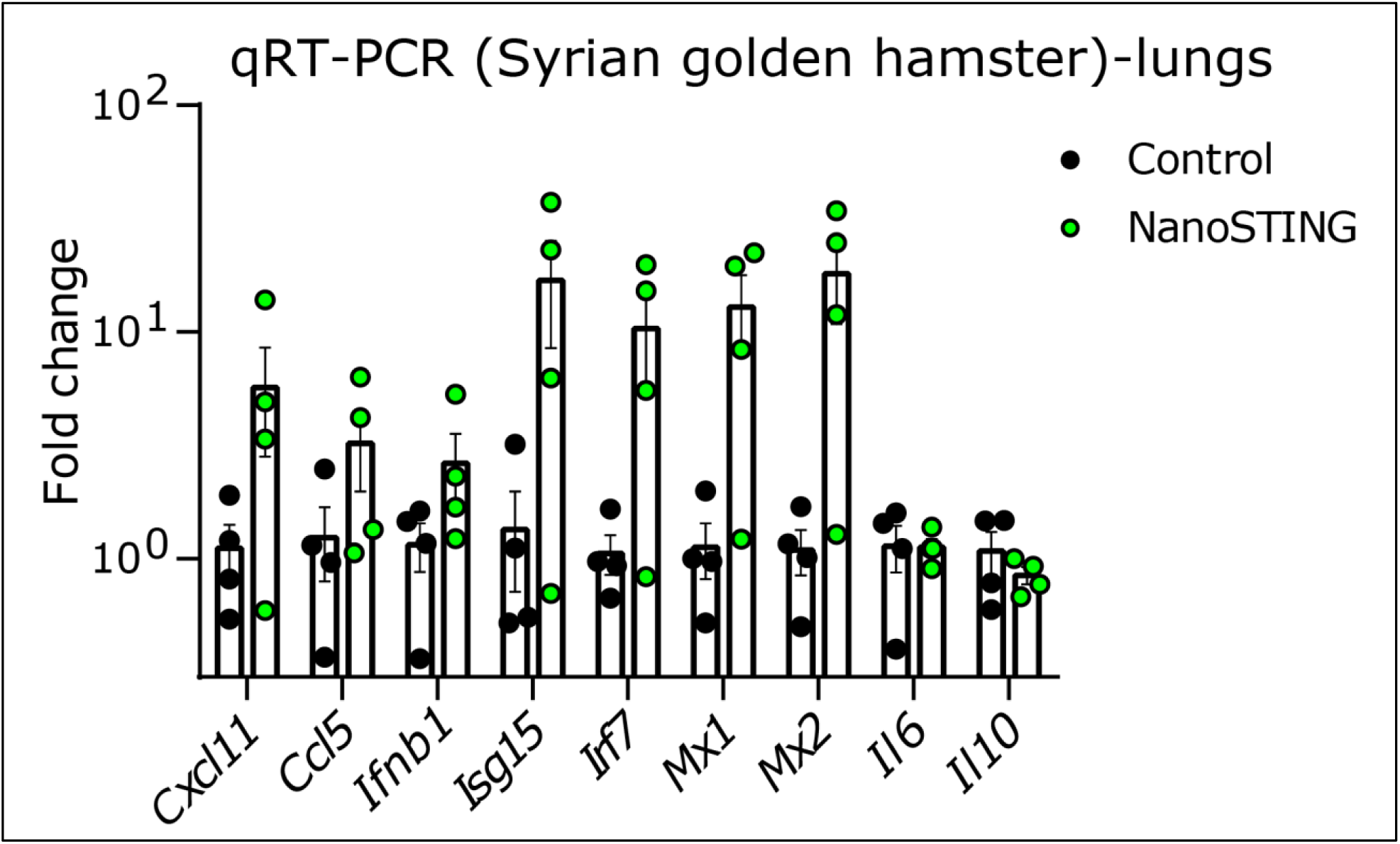
Upregulation of interferon-stimulated genes (*Cxcl11, Ccl5, Ifnb1, Isg15, Irf7, Mx1, Mx2, Il6,* and *Il10)* in lungs of hamsters post NanoSTING treatment (related to **Figure 3**) See Table S3 for a list of primers used. *For fold changes in gene expression, analysis was performed using a Mann-Whitney test. Vertical bars show mean values with error bar representing SEM. Each dot represents individual hamster*. *\*\*\*\*p < 0.0001; ***p < 0.001; **p < 0.01; *p < 0.05; ns, not significant*.

**Figure S6:**
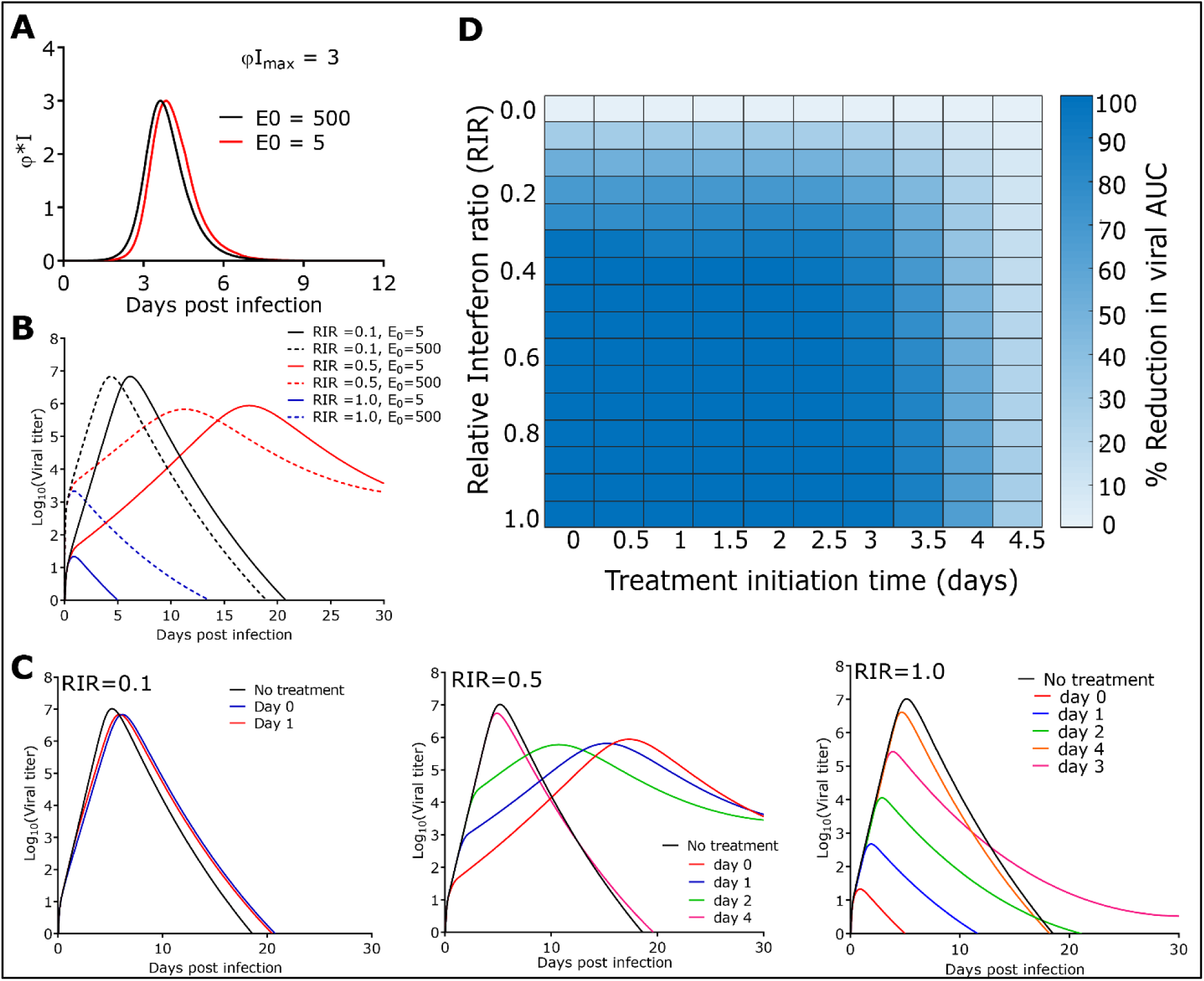
Sensitivity analysis (related to **Figure 4**) (A) Peak natural response is independent of the initial viral titer (B) Viral dynamics are independent of initial viral titer upon treatment with NanoSTING. E0 is the initial number of infected cells upon viral infection, which is a surrogate for viral titer. (C) Evolution of viral dynamics with different treatment initiation time and NanoSTING efficacies. (D) Heatmap of viral AUC with varying NanoSTING efficacy and treatment initiation time when NanoSTING effects last for 48 h after treatment initiation.

**Figure S7:**
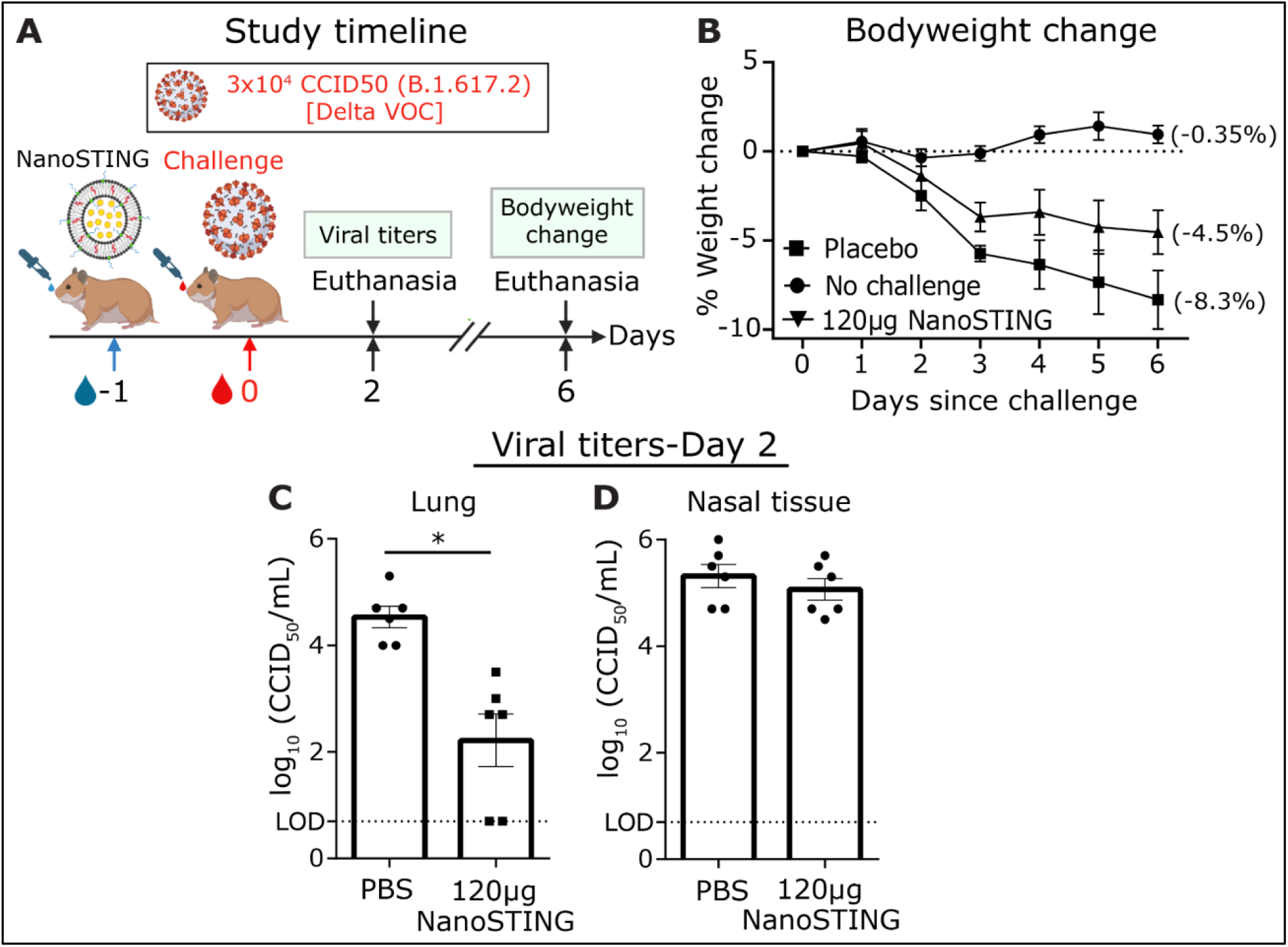
Pre-treatment of NanoSTING protects against challenge with SARS-CoV-2 delta VOC (B.1.617.2) (related to **Figure 5**) (A) Experimental setup. We treated groups of 6 hamsters, each with a single dose of NanoSTING (120 µg) and 72 h later challenged with ∼3 x 10^4^ CCID of SARS-CoV-2 virus (Delta VOC-B.1.617.2). We monitored animal weight changes daily for 6 days. Bodyweight changes and viral titers in the lungs and nasal tissue were used as primary endpoints. (B) Percentage of bodyweight change of NanoSTING treated animals compared to the control. (C, D) Viral titers were quantified in the lung and nasal tissue by plaque assay post day 2 after the challenge. The dotted line indicates limit of detection of the assay (LOD). *For viral titers, analysis was performed using a Mann-Whitney test. Vertical bars show mean values with error bar representing SEM. Each dot represents an individual hamster. Weight data was compared via mixed-effects model for repeated measures analysis. Lines depict group mean body weight change from day 0; error bars represent SEM. Asterisks indicate significance compared to the placebo-treated animals at each time point*. *\*\*\*\*p < 0.0001; ***p < 0.001; **p < 0.01; *p < 0.05; ns, not significant*.

**Figure S8:**
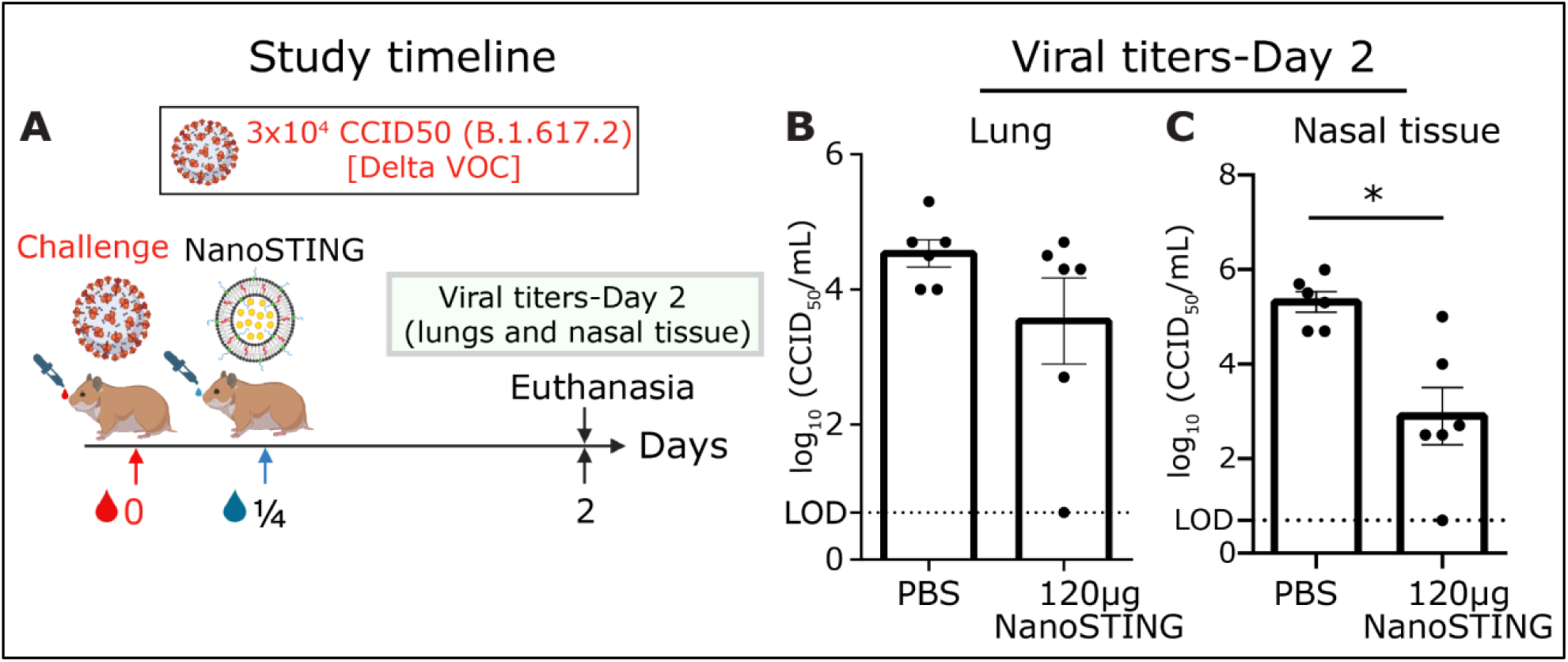
Post-treatment of NanoSTING protects against challenge with SARS-CoV2 delta VOC (B.1.617.2) (related to **Figure 5**) (A) Experimental setup. We challenged groups of 6 hamsters, each with ∼3 x 10^4^ SARS-CoV2 virus (Delta VOC-B.1.617) and 6 h later treated with a single dose of NanoSTING (120 µg). We euthanized the animals post day 2 of infection and determined viral titers in the lungs and nasal tissue. (B, C) Viral titers were quantified in the lung and nasal tissue by plaque assay post day 2 of infection. The dotted line indicates limit of detection of the assay (LOD). *For viral titers, analysis was performed using a Mann-Whitney test. Vertical bars show mean values with error bar representing SEM. Each dot represents an individual hamster. Asterisks indicate significance compared to the placebo-treated animals*. *\*\*\*\*p < 0.0001; ***p < 0.001; **p < 0.01; *p < 0.05; ns: not significant*.

**Figure S9:**
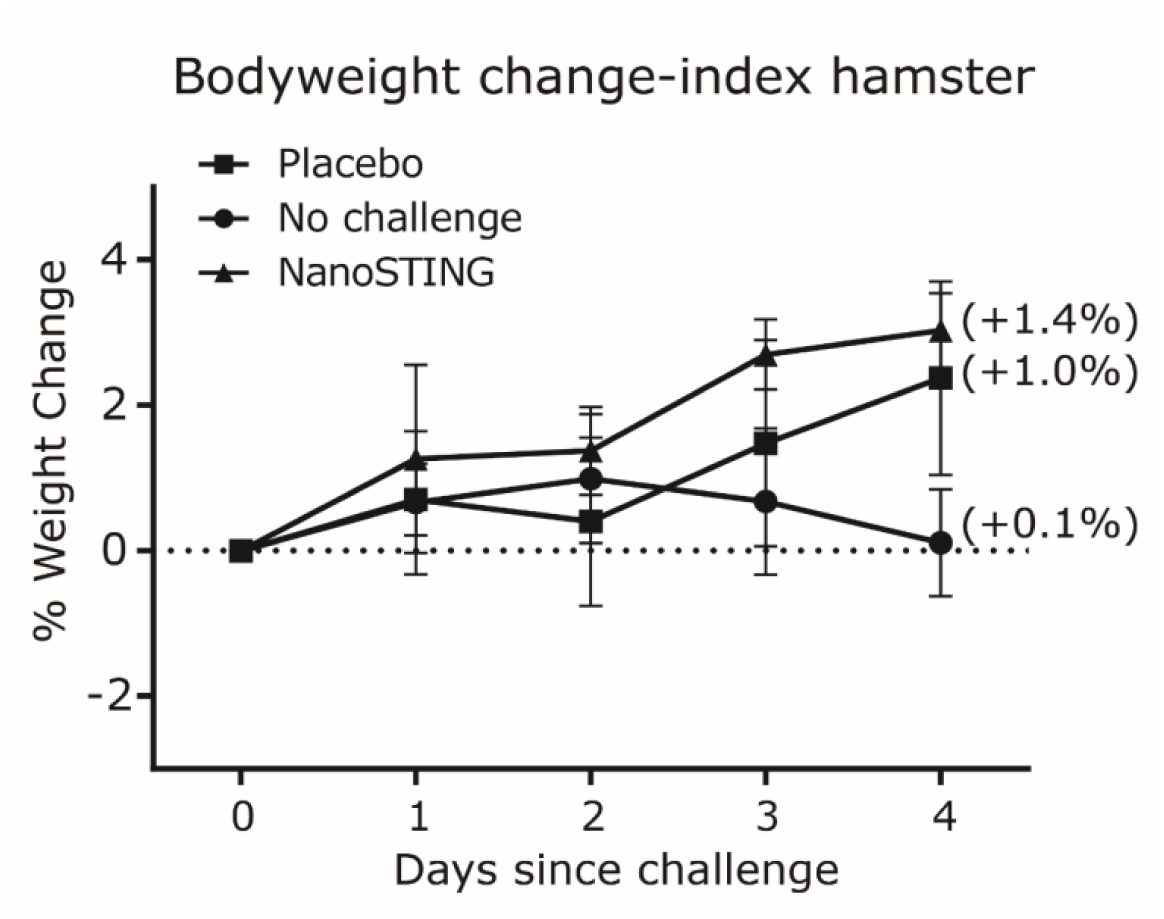
Longitudinal measurements of the bodyweight of index hamsters intranasally infected with SARS-CoV-2 Omicron VOC (related to **Figure 7**) *Weight data was compared via mixed-effects model for repeated measures analysis. Lines depict group mean body weight change from day 0; error bars represent SEM. Asterisks indicate significance compared to the placebo-treated animals at each time point*. *\*\*\*\*p < 0.0001; ***p < 0.001; **p < 0.01; *p < 0.05; ns: not significant*.

**Figure S10:**
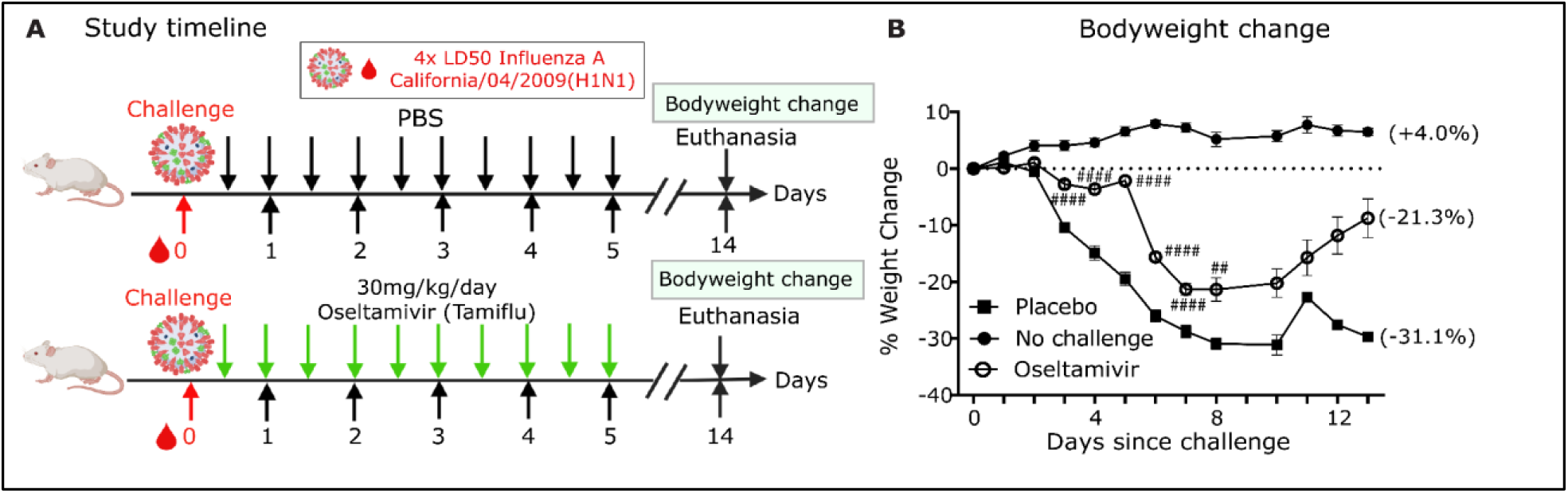
Treatment with Oseltamivir offers moderate protection against Influenza A (related to **Figure 8**) (A) Groups of ten BALB/c mice were challenged intranasally with ∼2 x 10^4^ of Influenza A/California/04/2009 (H1N1dpm) virus and treated with oseltamivir, twice daily, for five days. (B) Percent weight change of the different groups of mice. *Weight data was compared via mixed-effects model for repeated measures analysis. Lines depict group mean body weight change from day 0; error bars represent SEM. Pound sign show statistical significant differences between the Oseltamivir-treated group and placebo-treated animals*. *\*\*\*\*p < 0.0001; ***p < 0.001; **p < 0.01; *p < 0.05; ns, not significant*.

**Table S1.**
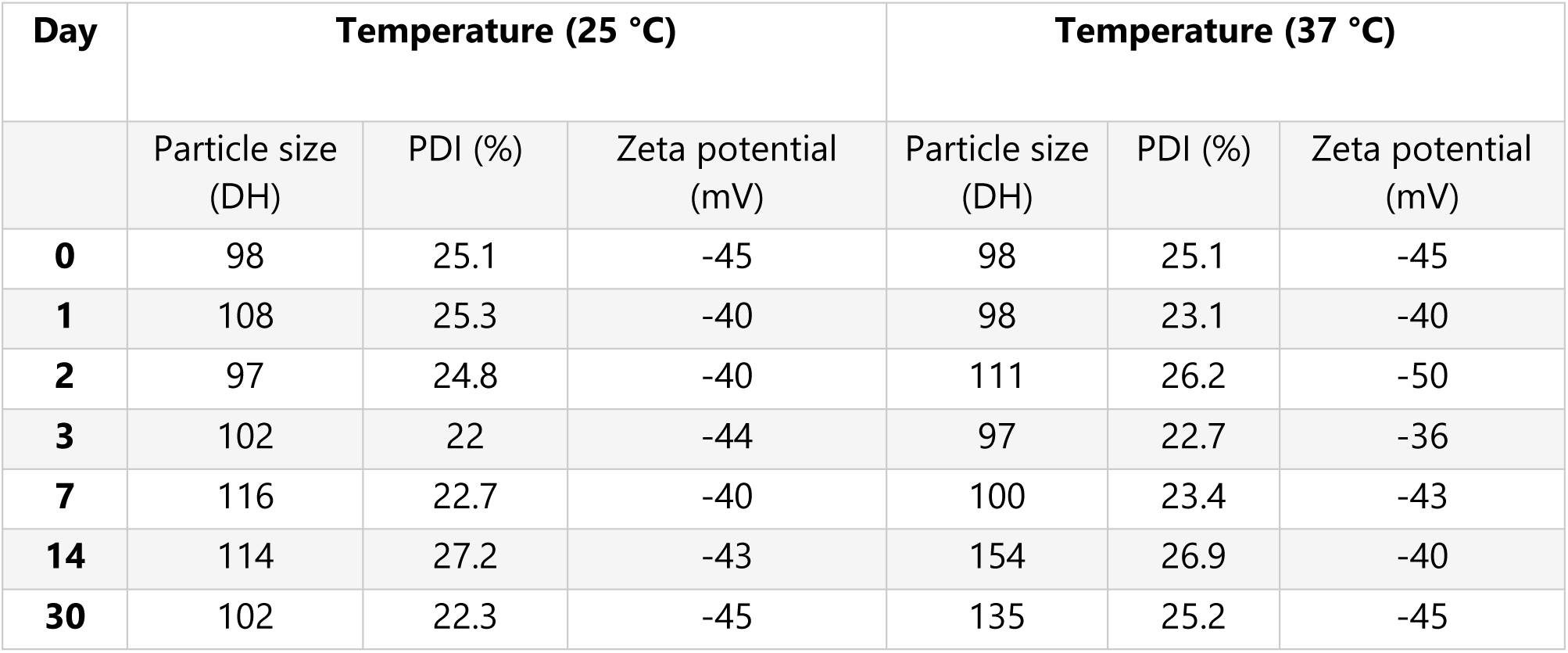
Stability studies of NanoSTING.

**Table S2.**
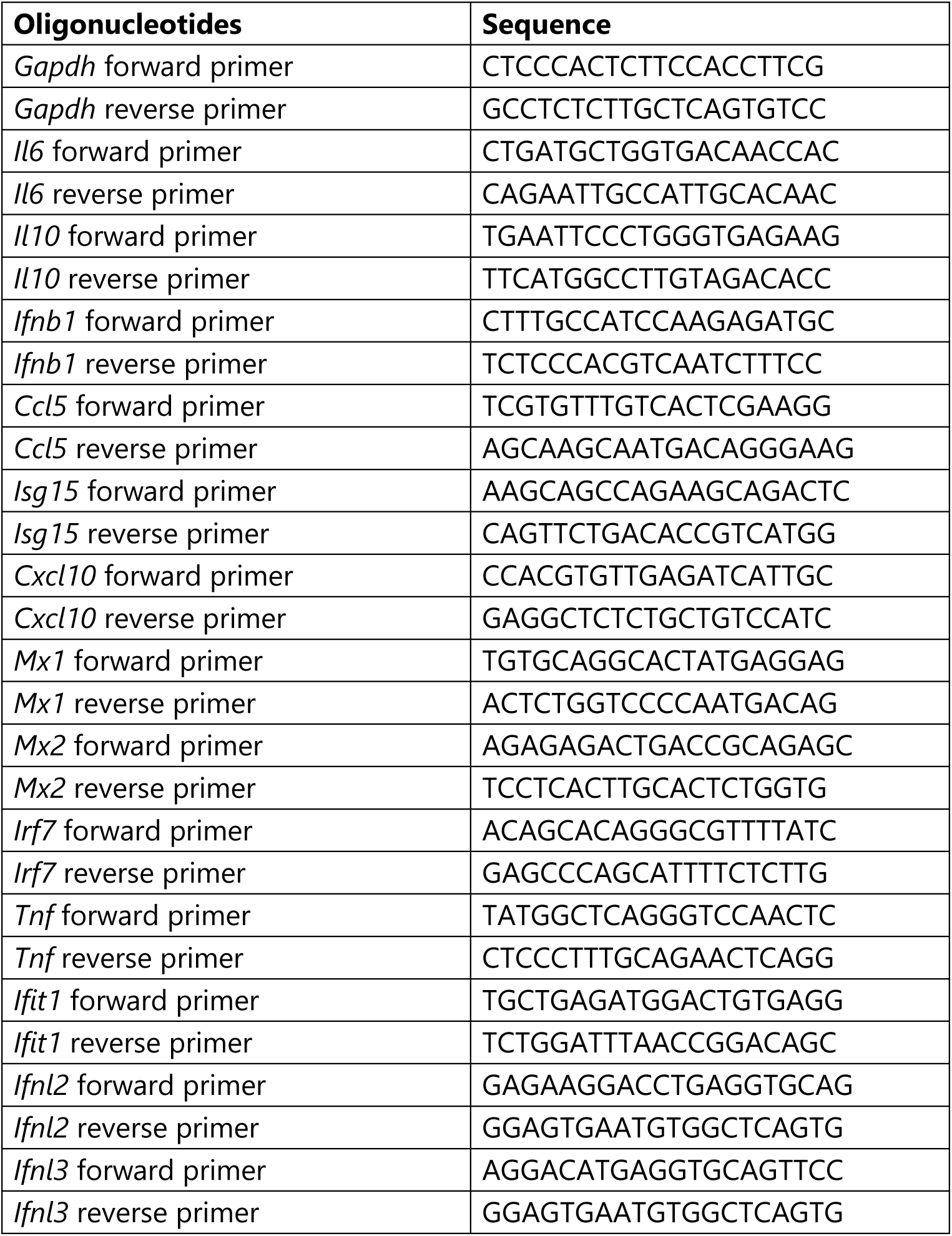
Primers used for qRT-PCR for Mus musculus BALB/c.

**Table S3.**
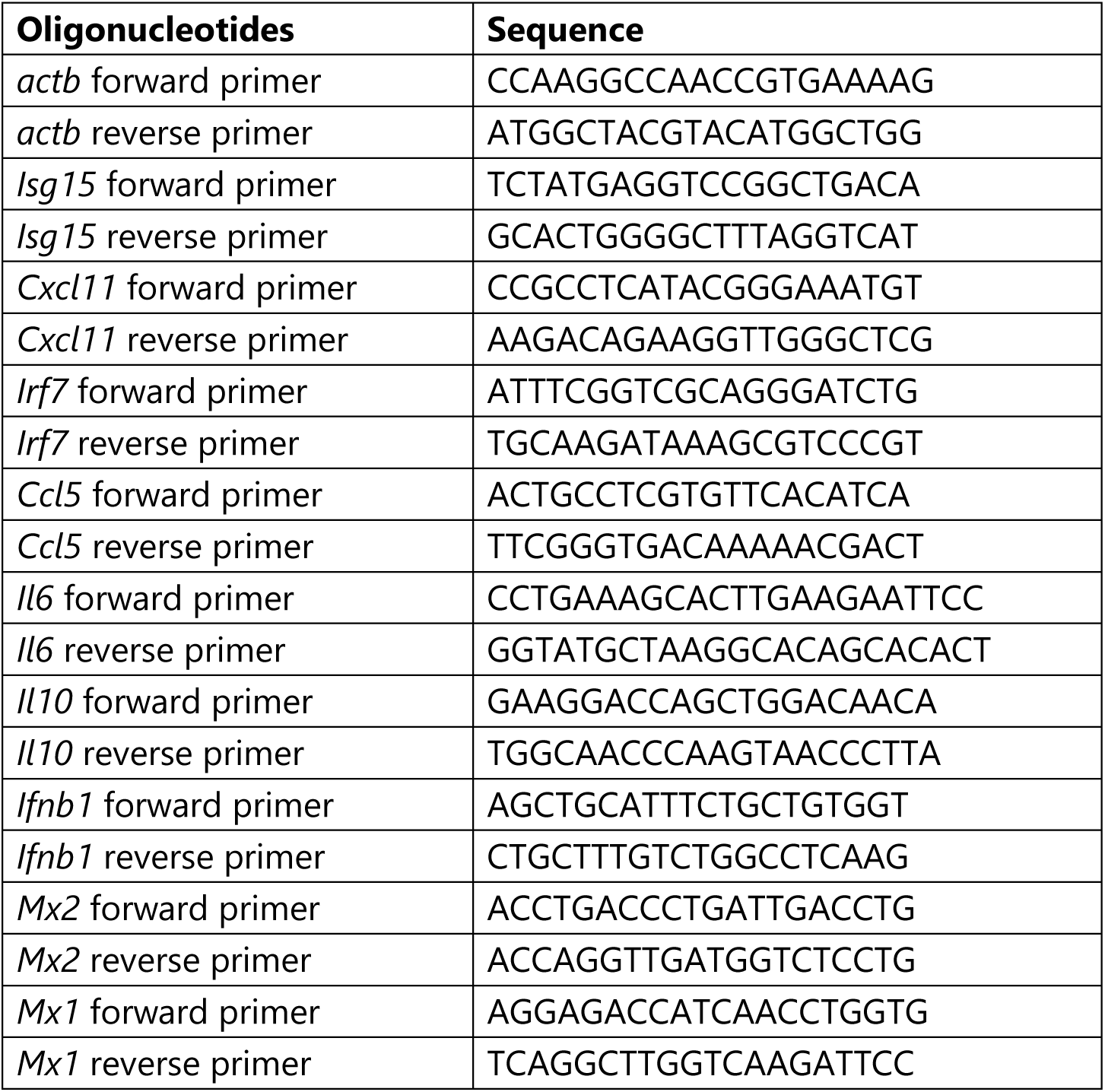
Primers used for qRT-PCR for *Mesocricetus auratus* (Syrian golden hamster)

## Supplementary methods

To quantify the kinetics of SARS-CoV-2 infection in the upper respiratory tract (URT) in the presence of NanoSTING, we used the Innate immune model described by Ke et al. ^34^. Assuming that NanoSTING efficacy is primarily due to the cell’s increased capacity to become refractory to infection, we modified the governing equation accordingly, as shown in the table below and Figure 4 of the main manuscript.

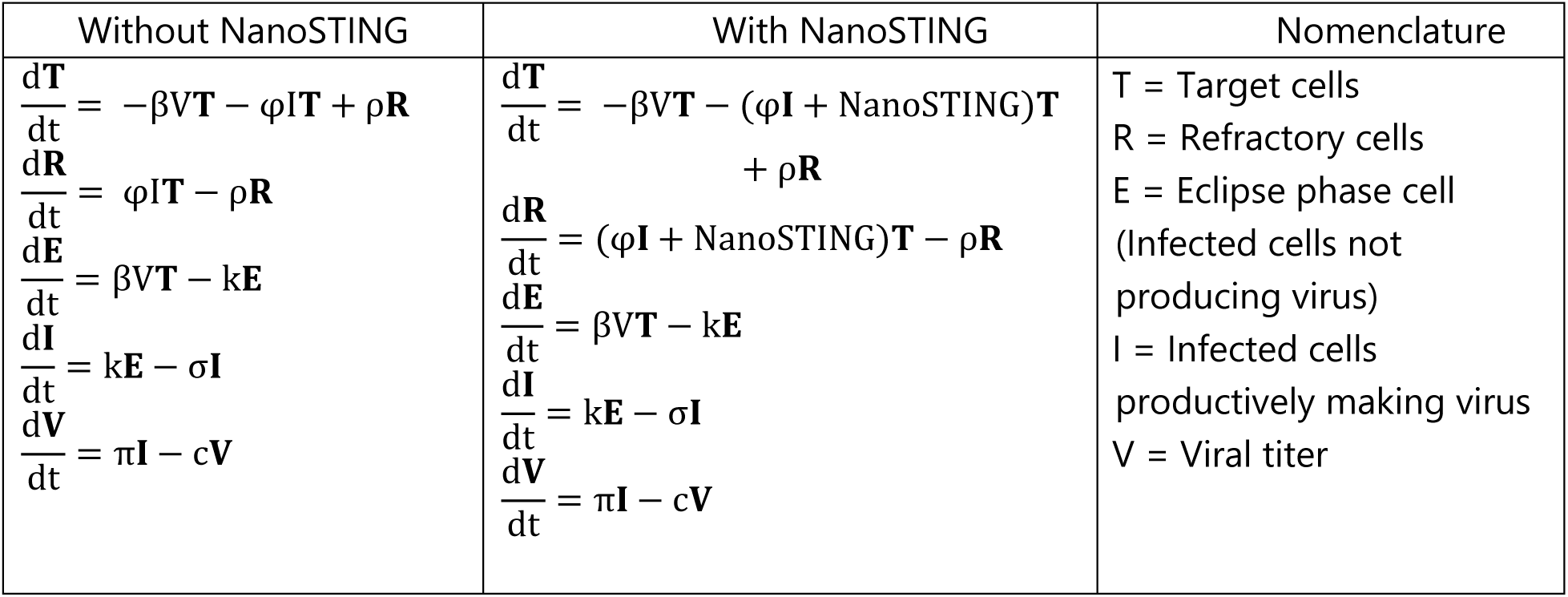

To get a physical interpretation of the variable NanoSTING, we non dimensionalized the target cell equation in the following way:

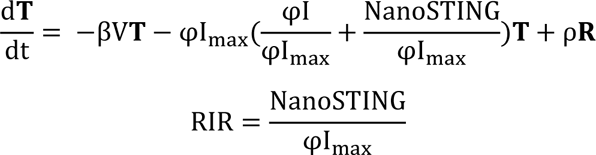

Where RIR is the relative interferon ratio, which is the relative contribution of NanoSTING to antiviral Interferon (refractory) responses compared to peak antiviral Interferon responses during SARS-CoV-2 without NanoSTING.

We solved these ordinary differential equations with mean population parameter values and initial values taken from Ke et. al^34^ and as shown in Table 1 & 2. First, we performed sensitivity analysis to show that the peak natural SARS-COV-2 response was independent of initial viral titer (Figure S6 A). We also performed sensitivity analysis to show that NanoSTING was effective at higher viral titers as well (Figure S6 B). We calculated the viral titer area under the curve (AUC) during infection for varying RIRs and the treatment initiation time post viral exposure. Because the effect of NanoSTING lasts only for 24 to 48 h, the NanoSTING coefficient was non zero only upto 24-48 h post treatment initiation.

**Table 1.**

| Parameter | Description | Mean population value |
| --- | --- | --- |
| $\beta$ | Infectivity parameter constant | $3.2 \cdot 10^{-8}$ mL/RNA copy/day |
| $\sigma$ | Death rate of infected cells | 1.7 /day |
| $\pi$ | Composite parameter for virus production and sampling | 45.3/mL/day |
| $\phi$ | Rate constant for Interferon induced conversion of Target cells to refractory cells | $1.3 \cdot 10^{-6}$ /cell/day |
| k | 1/the eclipse phase duration | 4 /day |
| c | virus clearance rate | 10 /day |
| $\rho$ | Rate at which refractory cells become target cells again | 0.0044/day |

**Table 2.**

| Variable | Initial value |
| --- | --- |
| T0 – Total number of target cells | $8 \cdot 10^7$ |
| E0 – Initial number of Infected cells | 5, 500 |

Sample MATLAB code:

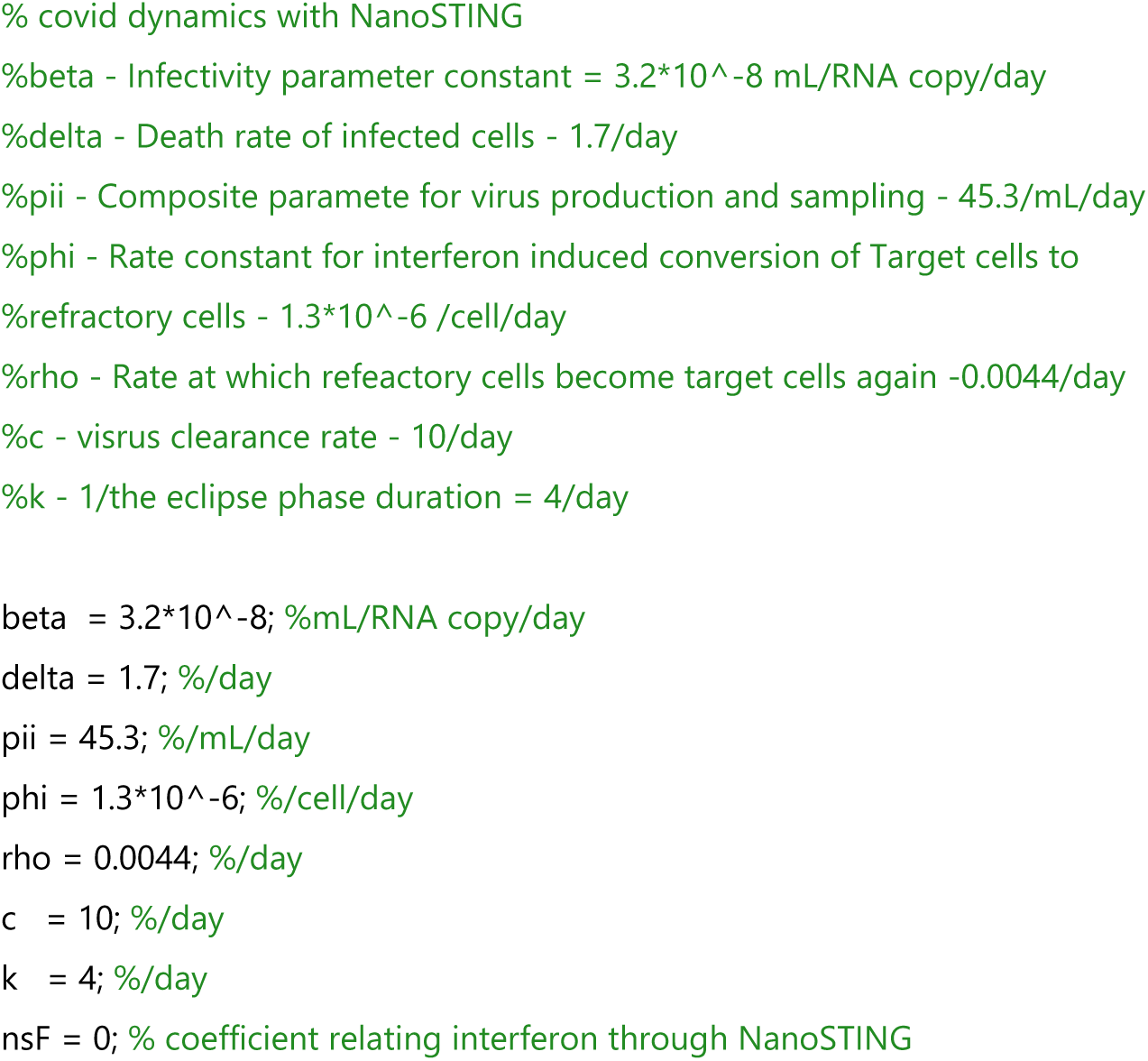
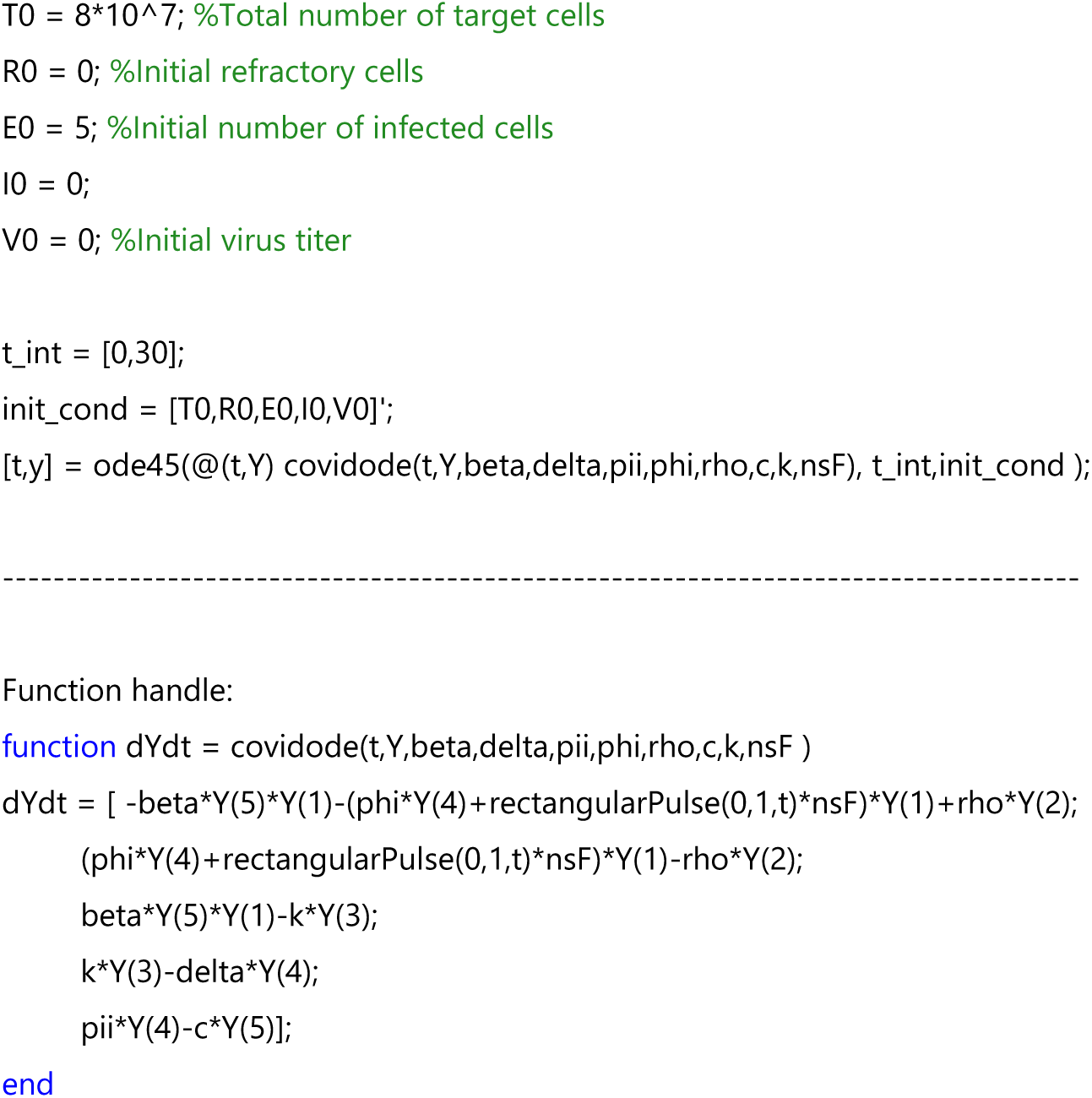

